# Haploinsufficiency of lysosomal enzyme genes in Alzheimer’s disease

**DOI:** 10.1101/2024.11.16.623962

**Authors:** Bruno A. Benitez, Clare E. Wallace, Maulikkumar Patel, Niko-Petteri Nykanen, Carla M. Yuede, Samantha L. Eaton, Cyril Pottier, Arda Cetin, Matthew Johnson, Mia T. Bevan, Woodrow D. Gardiner, Hannah M. Edwards, Brookelyn M. Doherty, Ryan T. Harrigan, Dominic Kurian, Thomas M. Wishart, Colin Smith, John R. Cirrito, Mark S. Sands

## Abstract

There is growing evidence suggesting that the lysosome or lysosome dysfunction is associated with Alzheimer’s disease (AD). Pathway analysis of post mortem brain-derived proteomic data from AD patients shows that the lysosomal system is perturbed relative to similarly aged unaffected controls. However, it is unclear if these changes contributed to the pathogenesis or are a response to the disease. Consistent with the hypothesis that lysosome dysfunction contributes to AD pathogenesis, whole genome sequencing data indicate that heterozygous pathogenic mutations and predicted protein-damaging variants in multiple lysosomal enzyme genes are enriched in AD patients compared to matched controls. Heterozygous loss-of-function mutations in the palmitoyl protein thioesterase-1 (*PPT1*), α-L-iduronidase (*IDUA*), β-glucuronidase (*GUSB*), N-acetylglucosaminidase (*NAGLU*), and galactocerebrosidase (*GALC*) genes have a gene-dosage effect on Aβ_40_ levels in brain interstitial fluid in C57BL/6 mice and significantly increase Aβ plaque formation in the 5xFAD mouse model of AD, thus providing *in vivo* validation of the human genetic data. A more detailed analysis of *PPT1* heterozygosity in 18-month-old mice revealed changes in α-, β-, and γ-secretases that favor an amyloidogenic pathway. Proteomic changes in brain tissue from aged *PPT1* heterozygous sheep are consistent with both the mouse data and the potential activation of AD pathways. Finally, CNS-directed, AAV-mediated gene therapy significantly decreased Aβ plaques, increased life span, and improved behavioral performance in 5xFAD/PPT1+/- mice. Collectively, these data strongly suggest that heterozygosity of multiple lysosomal enzyme genes represent risk factors for AD and may identify precise therapeutic targets for a subset of genetically-defined AD patients.

**Significance Statement:** Lysosomes play a role in the degradation of aggregation-prone proteins such as amyloid β (Aβ). Homozygous lysosomal enzyme gene defects result in fatal pediatric lysosomal storage diseases and, historically, carriers were considered normal. However, a human genetic analysis identified deleterious heterozygous variants in multiple lysosomal enzyme genes that are enriched in Alzheimer’s disease (AD) patients. Those findings were validated *in vivo* by demonstrating that heterozygous loss-of-function (LoF) mutations in five different lysosomal enzyme genes affect Aβ processing and exacerbate Aβ plaque formation. CNS-directed gene therapy ameliorated the effects of a heterozygous LoF mutation in one of those genes in a mouse model of AD. These findings provide insights into the role of lysosomes in AD and have important therapeutic implications.

## Introduction

Alzheimer’s disease (AD) is the most common form of dementia currently affecting ∼6.7 million people in the United States and 44 million worldwide, and these numbers are growing (1, 2). AD is characterized histologically by neuronal degeneration, extracellular β-amyloid (Aβ) plaques and intracellular Tau accumulation (1). Familial forms of AD are typically driven by increased Aβ production and subsequent aggregation of Aβ into soluble oligomers or insoluble extracellular plaques. However, studies in late-onset sporadic AD patients demonstrate impaired clearance of Aβ (3). Thus, the balance between production and clearance determines Aβ levels and the propensity to develop Aβ plaques.

The lysosome plays a major role in the degradation and clearance of intracellular organelles and aggregate-prone proteins (4). Genome wide association studies (GWAS) have consistently identified endo-lysosomal genes as being associated with sporadic and late-onset AD (5). Whole exome sequencing approaches have identified associations with coding variants in genes from the endolysosomal pathway in sporadic, early-onset AD (6). These genetic studies suggest that the lysosome may act as a common pathway through which downstream pathophysiological effects of AD might be mediated.

There is also a growing body of biological data supporting the role of lysosomes and lysosome dysfunction in AD. Lysosome biogenesis is up-regulated at early stages in the AD brain and AD models (7, 8). Lysosome dysfunction and β-amyloidogenesis are intertwined since lysosomes play a role in amyloid precursor protein (APP) processing. *Presenilin*1 (*PSEN1*) deletion elevates endolysosomal pH by disrupting vATPase assembly (9). The essential γ-secretase components, PSEN2 and Nicastrin, are located in the lysosome (10). Pharmacological impairment of lysosomal function *in vitro* results in changes in Aβ production (11, 12) and induces neuritic dystrophy in the absence of Aβ deposition *in vivo* (13). Changes in lysosomal pH reduce Aβ secretion (11) and lysosomal protease inhibitors reduce production of amyloidogenic Aβ peptides (12, 14). Homozygous loss-of-function mutations in several lysosomal enzyme genes result in the accumulation of Tau or altered Aβ metabolism and distribution (15–19). Accumulation of lysosomal markers are found in human brain tissue from patients with primary tauopathies (20). In the brains of patients with mucopolysaccharidosis (MPS) and Niemann-Pick type C (NPC) disease there is intense, intracellular Aβ staining throughout the brain (21, 22). MPS patients exhibit a significant increase in the level of soluble Aβ compared to normal control brains (22). Increased CSF levels of amyloidogenic Aβ peptides suggest an increased γ-secretase-dependent Aβ release in the brains of patients with NPC (23). In addition, mice deficient in *NPC1 (*Niemann-Pick C disease), *CLN3 (*Batten disease), *IDUA (*MPS-I), *SGSH (*Sanfilippo A), *GBA (*Gaucher disease), *TPP1 (*Late Infantile Batten disease) and *HEXB* (Sandhoff disease) genes exhibit an increase in both intracellular APP fragments (α-CTF/β- CTF) and Aβ_40/42_ levels (24–26). The absence of lysosomal neuraminidase 1 (*NEU1*) exacerbated the Aβ pathology in the 5XFAD mouse model of AD (17). Finally, selective enhancement of lysosomal proteolysis markedly reduces Aβ levels and amyloid load in AD mouse models (27, 28).

Although both genetic and biological studies support the role of lysosome dysfunction in AD pathogenesis, it is unclear what level of dysfunction is necessary to promote AD pathogenesis. Homozygous loss-of-function (LoF) mutations in lysosomal enzyme genes result in fatal pediatric disorders, collectively known as lysosomal storage diseases. Historically, carriers (heterozygotes) of LoF mutations in lysosomal enzyme genes were considered normal throughout life. In other words, 50% levels of lysosomal enzymes are believed to be sufficient for normal overall lysosome function and the prevention of disease. However, there are now several well-established examples of heterozygous deleterious mutations in lysosomal proteins that are associated with adult-onset neurological disease. The best-known example is heterozygosity of the glucocerebrosidase gene (*GBA*) and Parkinson disease (PD) (29). In fact, with the exception of age, heterozygosity of *GBA* is one of the greatest risk factors for PD. Similarly, heterozygous mutations in the progranulin gene (*GRN*) are associated with frontotemporal dementia (FTD) (30). Most recently, a severe heterozygous mutation in the Neimann-Pick C1 gene (*NPC1*) is associated with an apparently autosomal dominant, late-onset form of AD (31).

Based on strong evidence suggesting that the lysosome plays a critical role in Aβ metabolism and the fact that heterozygous mutations in several lysosomal proteins are associated with adult-onset neurological diseases, we hypothesized that heterozygous deleterious mutations in lysosomal enzyme genes are associated with AD. Here we provide human proteomic and genetic evidence supported by *in vivo* experimental data in both large and small animal models strongly suggesting that heterozygosity of multiple lysosomal enzyme genes is associated with AD. We further show that CNS-directed, AAV-mediated gene therapy can ameliorate some of the pathological and clinical signs in a mouse model of AD harboring a heterozygous LoF mutation in one of the genes. These data support the role of lysosome dysfunction in the pathogenesis of AD and identify potential therapeutic strategies for a subset of genetically-defined AD patients.

## Results

### Genetic and proteomic analyses in Alzheimer’s disease

In order to test the hypothesis that haploinsufficiency of lysosomal enzyme genes is associated with AD, coding variants in 44 lysosomal enzyme genes were identified using publicly available data bases. Known pathogenic mutations were identified in the ClinVar data base. The likelihood of variants of unknown significance (VUS) being deleterious was determined using the Combined Annotation Dependent Depletion (CADD) scoring system (32). Single-variant and gene-based analyses were performed on the 44 lysosomal genes in a case-controlled cohort of AD patients (Alzheimer’s Disease Sequencing Project, ADSP) containing 36,361 whole genome sequences. According to the genetic Principal Component Analyses, there were 6,547 and 4,739 Central European (CEU) AD patients and matched control samples, respectively. Similarly, 3,335 and 5,262 whole genome sequences were analyzed from African American (AA) AD patients and controls, respectively.

To investigate the impact of functional variants in endolysosomal genes, the analyses were restricted to ClinVar pathogenic variants and high and moderate variants with a CADD score greater than 30. In the CEU samples, SKAT-O analysis identified suggestive associations for the *PPT1* and *GAA* genes, and the Burden analysis identified a potential association for the *ASAH1* gene (Table 1). In the AA samples, potential associations were identified in the *IDUA* and *GALC* genes by both SKAT-O and Burden analyses (Table 1). The chromosomes, exact nucleotides, reference alleles, and alternate alleles for the *PPT1*, *GAA*, *GALC*, *IDUA*, and *ASAH1* genes are listed in Supplemental Table 1. By limiting a pathway level analysis to the ClinVar pathogenic variants in the 44 lysosomal enzyme genes, we identified a suggestive association (p = 8.63 x 10^-03^) with AD status using the SKAT-O analysis.

**Table 1:** Gene based analysis identified nominally significant associated genes in Central European* and African American^#^ populations in SKAT-O or Burden test analyses.

| Gene | Chromosome | Tested SNPs | C.MAF, Control | C.MAF, Cases | P-Value |
| --- | --- | --- | --- | --- | --- |
| * *GAA* <sup>1,£</sup> | 17 | 10 | $5.17 \times 10^{-03}$ | $7.25 \times 10^{-03}$ | $6.98 \times 10^{-03}$ |
| * *PPT1* <sup>1,£</sup> | 1 | 5 | 0 | $6.12 \times 10^{-04}$ | $3.83 \times 10^{-02}$ |
| #*GALC*<sup>1,2,€</sup> | 14 | 10 | $4.0 \times 10^{-04}$ | $2.30 \times 10^{-03}$ | $4.67 \times 10^{-02}$ |
| * *ASAH1* <sup>2,§</sup> | 8 | 3 | 0 | $3.16 \times 10^{-04}$ | $3.67 \times 10^{-02}$ |
| #*IDUA*<sup>1,2,£,€</sup> | 4 | 7 | $1.2 \times 10^{-02}$ | $1.9 \times 10^{-02}$ | $2.3 \times 10^{-02}$ |

C.MAF: Cumulative Minor Allele Frequency
<sup>1</sup>SKAT-O
<sup>2</sup>Burden Test
§ClinVar pathogenic variants
£CADD30 (High), ClinVar pathogenic variants
€CADD30 (High/Moderate), ClinVar pathogenic variants

Proteomic analysis was performed on tissue from the superior temporal gyrus (STG) from three late-stage patients and an equal number of controls that were ≥72 years of age in order to determine the effects of AD on the levels of various lysosomal proteins. The patients were diagnosed clinically and confirmed to have AD on post-mortem analysis (Supplemental Table 2, Supplemental Figure 1). The proteomic analysis was run in triplicate and 7,915 proteins were identified. The proteomic changes in the AD patients indicate convergence on the lysosomal storage disease pathway when interrogated using Ingenuity Pathway Analysis software (IPA, Supplemental Fig. 2). Thirty common lysosomal enzymes were identified, greater than half of which were differentially expressed by ≥15%, and of those 11 were increased by ≥20% (Table 2).

**Table 2:**
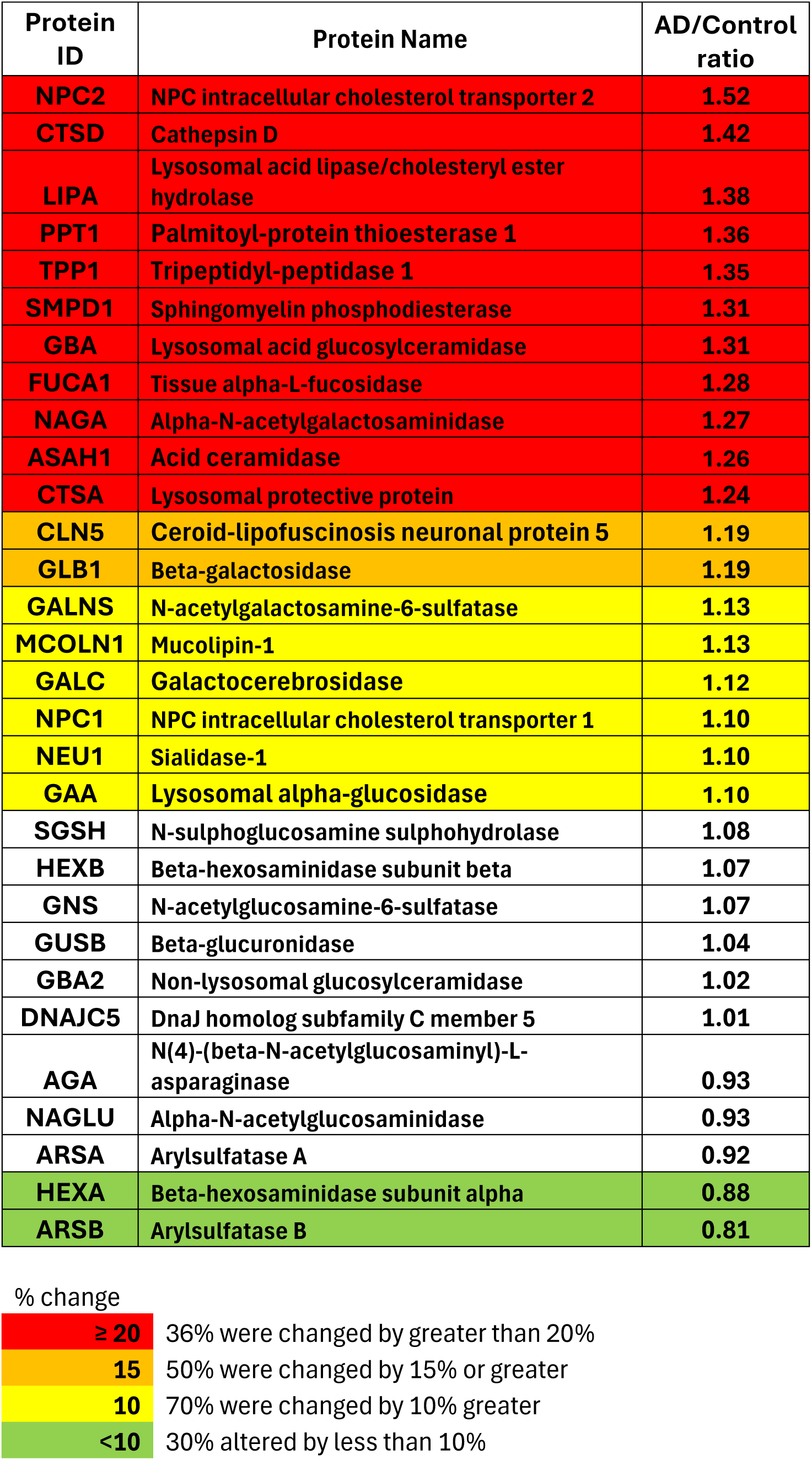
Changes in the levels of lysosomal proteins in brain tissue from AD patients compared to similarly aged controls.

### Haploinsufficiency of palmitoyl protein thioesterase-1 and Alzheimer’s disease

The human genetic and proteomic analyses support the hypothesis that lysosome dysfunction is associated with AD. In addition, they serve as an effective means to prioritize the lysosomal enzyme genes for further *in vivo* validation. Pathogenic mutations and deleterious variants in the *PPT1* gene are enriched in CEU AD patients. There are both large (ovine) (33) and small (murine) (34) animal models of CLN1 disease (infantile neuronal ceroid lipofuscinosis) which is caused by a complete deficiency of the lysosomal enzyme, palmitoyl protein thioesterase-1 (PPT1). In addition, several critical proteins responsible for Aβ metabolism are palmitoylated and changes in palmitoylation have been reported in AD (35). Therefore, we determined the effects of *PPT1* heterozygosity on the sheep brain proteome, in a mouse model of AD, and on Aβ metabolism in otherwise normal mice.

Samples from the entorhinal cortex from four *PPT1* heterozygous sheep (554-2022 days of age) and five normal controls (500-1335 days of age) were collected and processed for proteomic analysis (Supplemental Table 3). All samples were run in triplicate and 8,835 proteins were detected in the *PPT1* heterozygous sheep. Pathway analyses of data from the *PPT1* heterozygous sheep and controls indicate that multiple protein differences are associated with interactions known to play a role in AD (Supplemental Fig. 3). When the proteomic data from the heterozygous *PPT1* sheep are compared to the proteomic changes in AD patients, several important proteins in the amyloid processing pathway network also appear to be altered in a similar manner

To more precisely determine whether PPT1 haploinsufficiency exacerbates an amyloidogenic process *in vivo*, a *PPT1* null allele was bred onto the 5xFAD murine model of AD (36). The resulting animals were heterozygous for the *PPT1* gene on the 5xFAD background and are hereafter referred to as 5xFAD/PPT1+/- mice. These mice have approximately 50% normal levels of PPT1 activity in the brain (Fig. 1A) and appear to develop normally. The 5xFAD/PPT1+/- mice exhibit a significantly shortened life span (median ∼10 months) compared to either the 5xFAD or *PPT1* heterozygous mice (median ∼24 months) (Fig. 1B). Coronal sections of the brains from seven-month-old 5xFAD/PPT1+/- mice were analyzed for amyloid plaque burden using anti-amyloid antibodies (HJ3.4). There is an obvious (Fig. 1C) and statistically significant increase (Fig. 1D) in Aβ plaques in the 5xFAD/PPT1+/- mice compared to the parental 5xFAD strain. To complement the finding of increased in Aβ plaques caused by PPT1 haploinsufficiency, samples from the hippocampus of those animals were fractionated into detergent-soluble and guanidine-soluble (detergent-insoluble) fractions, and the Aβ isoforms were quantified by enzyme-linked immunosorbent assay (ELISA). The levels of detergent-insoluble Aβ_40_ and Aβ_42_ were significantly elevated in 5xFAD/PPT1+/- animals compared with 5xFAD (Fig. 1E, F).

**Fig. 1.**
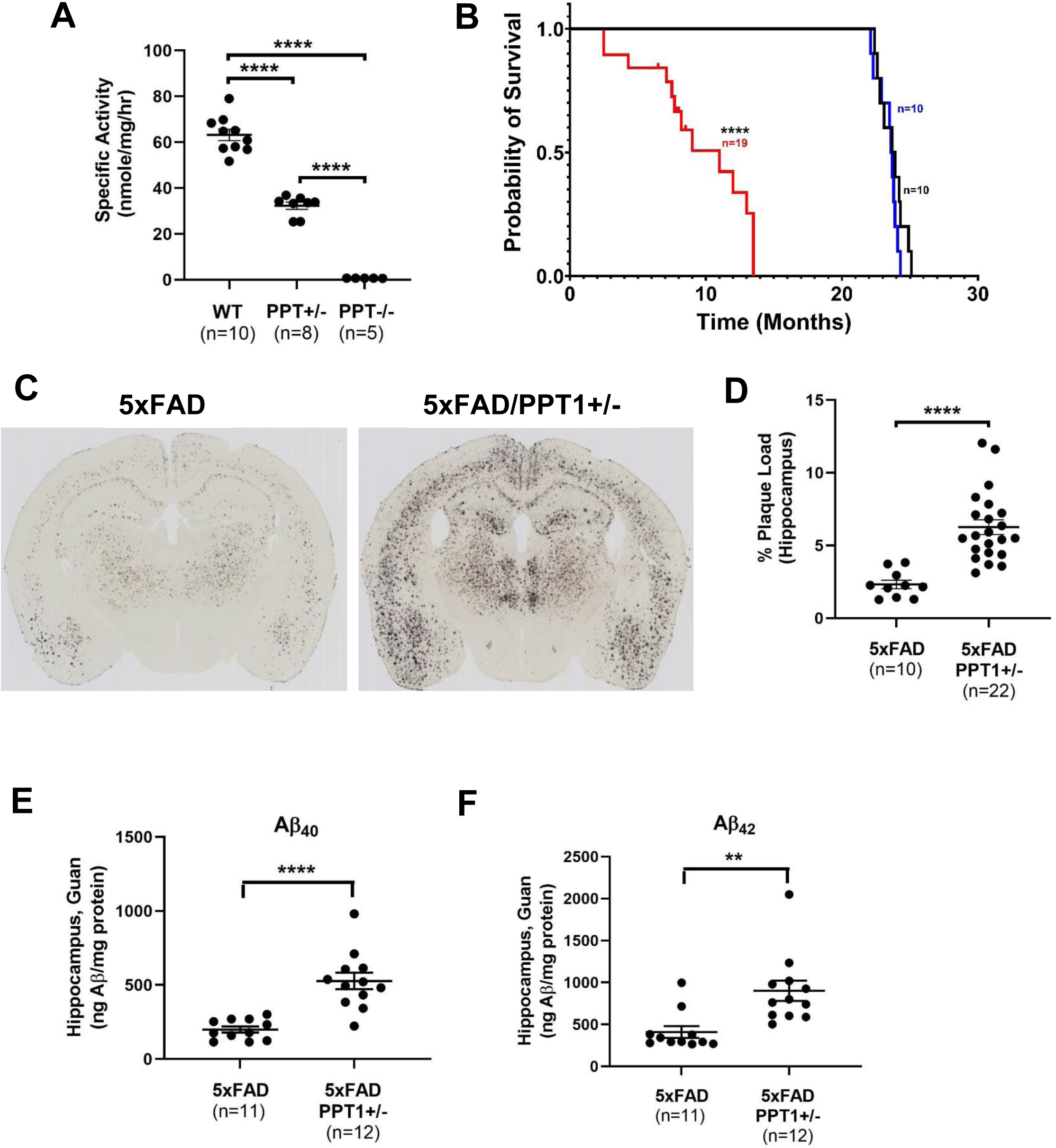
Heterozygosity of *PPT1* exacerbates the AD-like phenotype of the 5xFAD mouse. (A) Heterozygosity of *PPT1* decreases brain PPT1 activity by approximately half and homozygous deficiency completely eliminates enzyme activity. (B) The median life span of 5xFAD (blue line) and *PPT1* heterozygous mice (black line) on a C57Bl/6 background is 24 months. In contrast, the median life span of 5xFAD/PPT1+/- mice (red line) is approximately 10 months. (C) There is an obvious and statistically significant (D) increase in Aβ plaques in seven-month-old 5xFAD/PPT1+/- mice compared to age-matched 5xFAD mice. The levels of detergent-insoluble Aβ_40_ (E) and Aβ_42_ (F) in 5xFAF/PPT1+/- mice are significantly greater than those in age-matched 5xFAD mice. **p<0.01, ****p<0.0001.

### Haploinsufficiency of PPT1 affects Aβ metabolism

Although heterozygosity of *PPT1* exacerbates the phenotype of 5xFAD mice, it is important to determine the effects of haploinsufficiency in the absence of any transgenes that might otherwise affect Aβ metabolism. It has been demonstrated that the level of soluble Aβ in brain interstitial fluid (ISF) reflects the total soluble Aβ in extracellular pools (37). Therefore, the levels of murine Aβ in brain ISF were determined in the hippocampus of three-to four-month-old WT, heterozygous, and homozygous deficient *PPT1* animals on a C57Bl/6 background. There is a clear gene-dosage effect of *PPT1* on baseline Aβ levels in hippocampal ISF (Fig. 2A), suggesting that dysfunctional lysosomes are likely to prevent Aβ from being secreted from neurons.

**Fig. 2.**
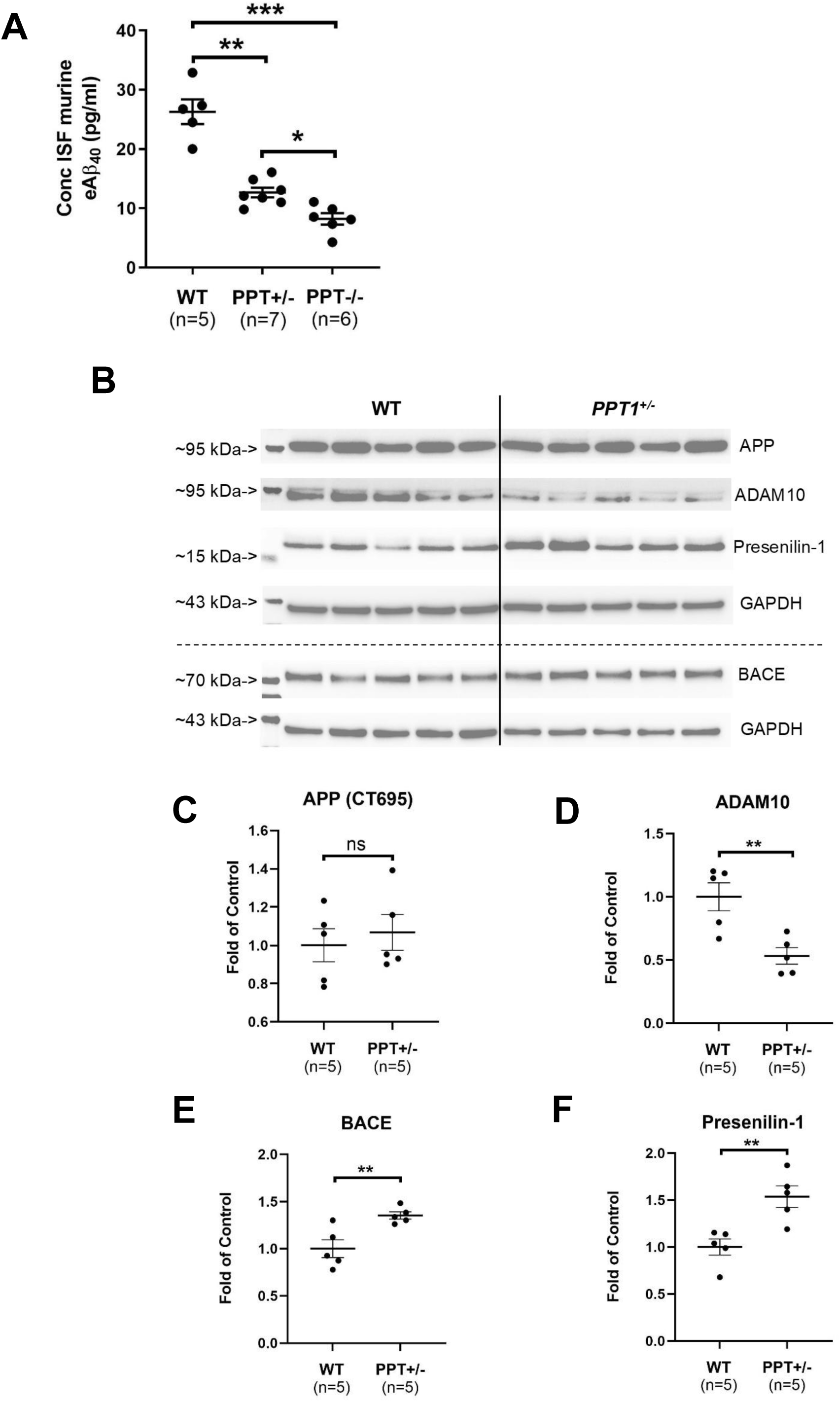
Heterozygosity of *PPT1*, alone, affects Aβ processing and the processing enzymes. There is a clear gene-dosage effect of PPT1 on ISF Aβ levels. (A) The level of ISF Aβ_40_ is significantly decreased in 3.5- month-old *PPT1* heterozygous animals compared to age-matched wild type animals. The level of ISF Aβ_40_ is further decreased in *PPT1* homozygous deficient mice compared to heterozygous animals. (B) Quantitative Western blots of APP, ADAM 10, Presenilin-1, and BACE are shown [Note: a single Western blot was probed with APP, ADAM10, Presenilin-1, and GAPDH antibodies; a second Western blot (below the dashed line) was probed with BACE and GAPDH antibodies]. (C) The levels of amyloid precursor protein [APP (CT695)] in 18- month-old *PPT1* heterozygous animals are not significantly different from age matched wild type mice. (D) The levels of α-secretase (ADAM10) are significantly decreased in PPT1 heterozygous mice. In contrast, the levels of both β-secretase (E) and γ-secretase (F) are significantly increased in *PPT1* heterozygous animals compared to wild type mice. *p<0.05, **p<0.01, ***p<0.001.

Western blot analysis was performed on brain homogenates from 18-month-old *PPT1* heterozygous mice and age-matched wild type (WT) animals on the C57Bl/6 background (Fig. 2B). The level of amyloid precursor protein (APP) in *PPT1* heterozygous mice is not significantly different from WT animals (Fig. 2C). The level of ADAM10 (α-secretase), which favors a non-amyloidogenic pathway, is significantly decreased in *PPT1* heterozygous mice (Fig. 2D). In contrast, the levels of β-secretase (BACE) (Fig. 2E) and the catalytic component of γ-secretase (presenilin-1) (Fig. 2F), which favor an amyloidogenic pathway, are significantly elevated in the 18-month-old heterozygous *PPT1* animals compared to the WT controls. Several critical proteins in the amyloid processing pathway network are changed in the brains of AD patients and *PPT1* heterozygous sheep. Interestingly, similar to the heterozygous *PPT1* mice, amyloid precursor protein is not elevated in the heterozygous sheep even though the AD pathway appears to be activated. Collectively, these data suggest that haploinsufficiency of PPT1 alone favors an amyloidogenic program.

### AAV-mediated gene therapy improves the phenotype of 5xFAD/PPT1+/- mice

CNS-directed, AAV-mediated gene therapy was performed in 5xFAD/PPT1+/- mice with an AAV9-hPPT1 gene transfer vector in order to determine if complementing the haploinsufficient levels of PPT1 could improve the exacerbated AD-like phenotype. We first determined if Aβ plaques could be reduced if the AAV vector was delivered during the neonatal period. Animals received four bilateral intraparenchymal injections of the AAV vector at post-natal day 1 or 2 to target the anterior cortex and hippocampus. At seven-months-of-age, there was a significant decrease in Aβ plaques in the AAV-treated animals compared to age-matched untreated 5xFAD/PPT1+/- mice (Fig. 3A). The level of PPT1 enzyme activity approached normal levels in the AAV-treated animals compared to the untreated controls (Fig. 3B).

**Fig. 3.**
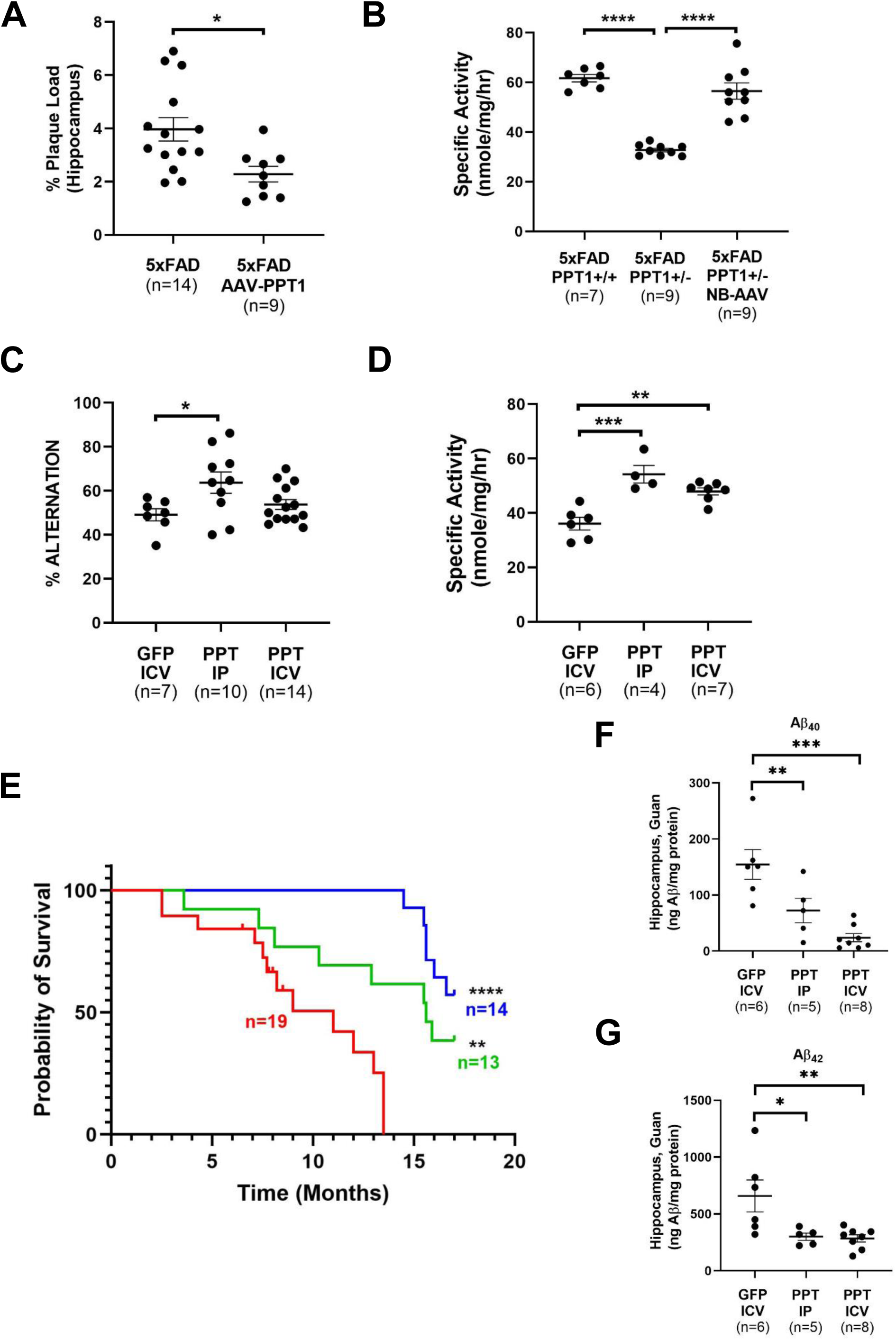
CNS-directed, AAV-mediated gene therapy improves the AD phenotype in 5xFAD/PPT1+/- mice. (A) Intracranial injection of an AAV9-hPPT1 vector in newborn (PND 1-2) 5xFAD/PPT1+/- mice significantly decreases the Aβ plaque load at 7 months-of-age compared to uninjected 5xFAD/PPT1+/- mice. (B) The level of PPT1 activity in AAV-hPPT1-injected mice is approximately 2-fold greater than uninjected mice. (C) 5xFAD/PPT1+/- mice receiving intraparenchymal injections of AAV-hPPT1 (PPT, IP) at 3.5-months of age performed significantly better in the Y-maze behavioral test compared to 5xFAD/PPT1+/- mice receiving an ICV injection of AAV9-GFP (GFP, ICV) at the same age. Although, on average, mice receiving an ICV injection of AAV9-hPPT1 (PPT, ICV) performed better than AAV9-GFP-injected animals, it did not reach statistical significance. (D) The level of PPT1 activity in 17-month-old 5xFAD/PPT1+/- mice receiving an ICV or an intraparenchymal injection was significantly greater than 5xFAD/PPT1+/- mice injected with AAV9-GFP. (E) 5xFAD/PPT1+/- mice receiving intraparenchymal injections of AAV-hPPT1 (green line) at 3.5-months of age had a significantly extended life span compared to untreated 5xFAD/PPT1+/- animals (red line). The life span of 5xFAD/PPT1+/- mice receiving an ICV injection of AAV9-hPPT1 (blue line) was significantly greater than untreated mice, and mice receiving an intraparenchymal injection of virus. The levels of detergent-insoluble Aβ_40_ (F) and Aβ_42_ (G) were significantly decreased in 17 month-old 5xFAD/PPT1+/- mice injected with AAV9-hPPT1 by either route compared with 9 month-old 5xFAD/PPT1+/- mice injected ICV with AAV9-GFP. *p<0.05, **p<0.01, ***p<0.001, ****p<0.0001.

The effect of delaying the AAV injections and administering them through different routes was then determined. Two groups of 5xFAD/PPT1+/- animals were treated at the same age (3.5 months) with the identical total dose (8 x 10^9^ vg) of vector. One group received a single intracerebroventricular (ICV) injection. The second group received the identical dose divided equally between four different intraparenchymal injection sites to target the anterior cortex and hippocampus. A third group received a single ICV injection (8 x 10^9^ vg) of a nearly identical AAV9 vector expressing GFP. At seven-months-of-age, the Y-maze and elevated plus maze behavioral tests were performed on those animals. The animals that received the intraparenchymal injections of AAV9- hPPT1 performed significantly better than AAV9-GFP-injected mice on the Y-maze (Fig. 3C). Although the mean performance of the animals receiving the ICV injections was improved compared to the AAV9-GFP-injected animals, it did not reach statistical significance. Similarly, although the mean performance of both groups receiving AAV9-hPPT1 in the elevated plus maze was improved compared to the AAV9-GFP-treated group, neither group reached statistical significance (Supplemental Fig. 4).

The AAV9-GFP-injected animals were sacrificed at nine months of age for biochemical analysis and the AAV9-hPPT1-injected animals were used for both life span determinations and biochemical analysis. The level of PPT1 activity in both groups of AAV9-hPPT1-treated animals at 17 months was significantly greater than 5xFAD/PPT1+/- animals receiving ICV injections of AAV9-GFP (Fig. 3D). Both groups of animals receiving injections of AAV9-hPPT1 had significantly increased life spans compared to untreated 5xFAD/PPT1+/- mice (Fig. 3E). Animals that received ICV injections of vector had a significantly increased life span compared to animals receiving intraparenchymal injections. At 17 months of age, 57% and 38% of the ICV- and intraparenchymal-injected mice, respectively, were still alive but were sacrificed for further analysis. Both groups of AAV9-hPPT1-treated mice had significantly reduced levels of detergent-insoluble Aβ_40_ and Aβ_42_ compared to younger (9 mo) AAV9-GFP-injected mice (Fig. 3F, G).

### Validation of additional genes identified in the human genetic analyses

Two additional genes that were identified in the genetic analysis were validated *in vivo*. Deleterious mutations/variants in the α-L-iduronidase (*IDUA*) gene were significantly enriched in the AA AD samples compared to controls (Table 1). Haploinsufficiency of IDUA in C57Bl/6 mice decreased enzyme activity (Fig. 4A) and resulted in a gene-dosage effect on murine Aβ_40_ levels in brain ISF compared to age-matched WT controls (Fig. 4B). When a single *IDUA* null allele was bred onto the 5xFAD mouse model of AD, both the plaque burden (Fig. 4C, Supplemental Fig. 5A) and level of insoluble Aβ_42_ (Fig. 4E) were significantly increased compared to 5xFAD mice. Although the mean level of insoluble Aβ_40_ was greater than that observed in the 5xFAD mice, it did not reach statistical significance (Fig. 4D).

**Fig. 4.**
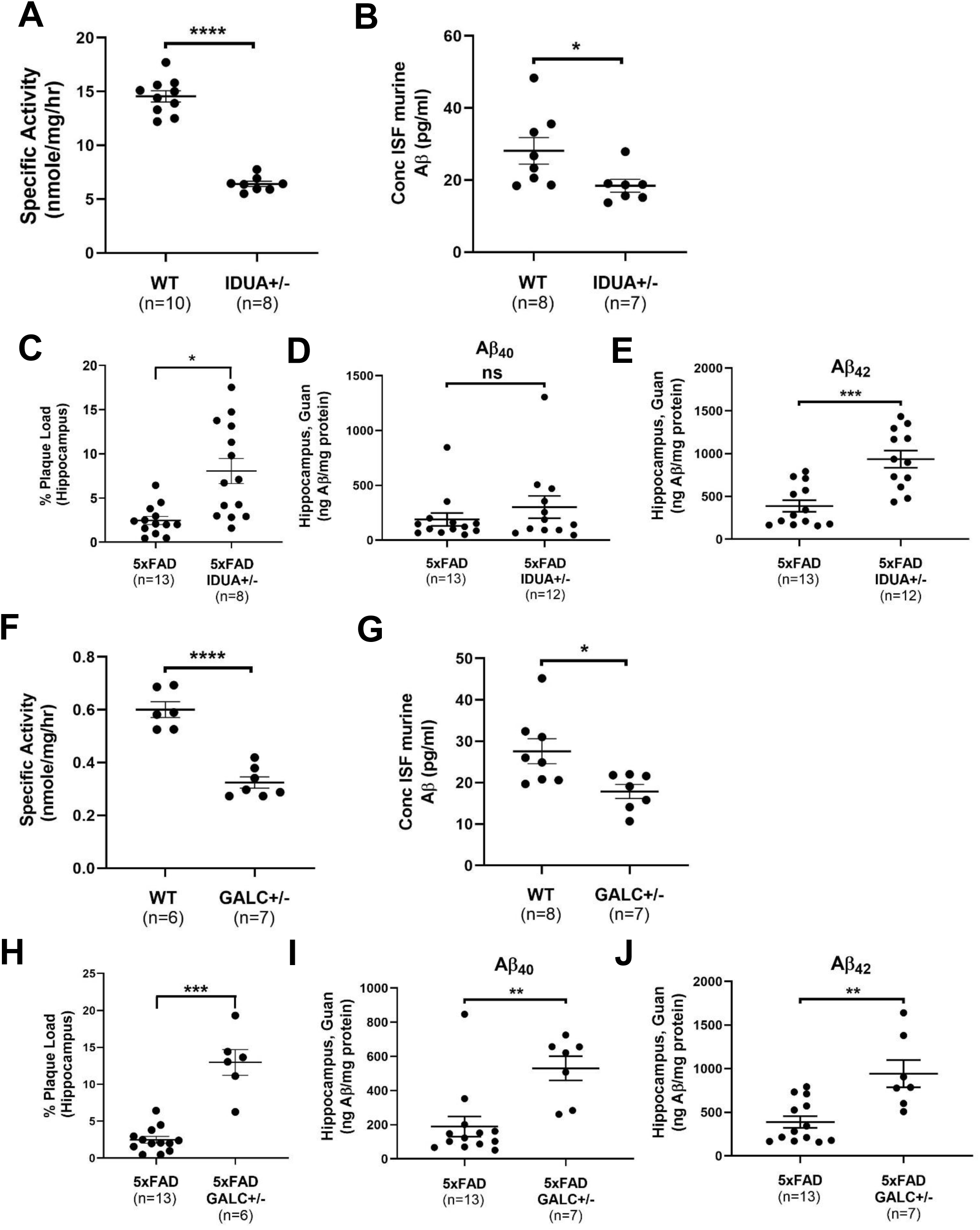
Two additional lysosomal enzyme genes identified in the human genetic analysis affect Aβ metabolism and exacerbate the AD phenotype of 5xFAD mice. Three and one half-month-old mice that are heterozygous for the α-L-iduronidase (*IDUA*) gene (IDUA+/-) have approximately 50% enzyme activity (A) and significantly decreased ISF Aβ levels (B) compared to age-matched wild type (WT) mice. Heterozygosity of *IDUA* also significantly increased the Aβ plaque load (C) and the level of detergent-insoluble Aβ_42_ in 5xFAD mice (E). Although the mean level of detergent-insoluble Aβ_40_ was greater than that observed in 5xFAD mice, it did not reach statistical significance (D). Three and one half-month-old mice that are heterozygous for the galactocerebrosidase (*GALC)* gene (GALC+/-) have approximately 50% enzyme activity (F) and significantly decreased ISF Aβ levels (G) compared to age-matched wild type mice. Heterozygosity of *GALC* also significantly increased the Aβ plaque load (H) and the level of detergent-insoluble Aβ_40_ (I) and Aβ_42_ (J) in 5xFAD mice (I, J). Note: The ELISA assays and Aβ immunostaining for the 5xFAD, 5xFAD/IDUA+/-, 5xFAD/GALC+/-, and 5xFAD/GUSB+/- animals were performed at the same time with the same reagents to ensure consistency for direct comparisons. Therefore, the values for the 5xFAD/IDUA+/-, 5xFAD/GALC+/-, and 5xFAD/GUSB+/- animals were compared to the same set of data from the 5xFAD mice (see Figures 4C, D, E, H, I, J, 5C, D, and E). *p<0.05, **p<0.01, ***p<0.001, ****p<0.0001.

Deleterious mutations/variants in the galactocerebrosidase (*GALC*) gene were also significantly enriched in the AA AD samples compared to controls (Table 1). Similar to the *PPT1* and *IDUA* genes, a heterozygous null allele of *GALC* decreased GALC enzyme activity (Fig. 4F) and had a gene-dosage effect on ISF Aβ levels in C57Bl/6 mice (Fig. 4G). Haploinsufficiency of GALC also increased the Aβ plaque load (Fig. 4H, Supplemental Fig. 5B) and insoluble Aβ_40_ and Aβ_42_ levels in the 5xFAD mouse (Fig. 4I, J).

### Lysosomal enzyme genes <u>not</u> identified in the human genetic analysis

The fact that the human genetic analyses identified multiple lysosomal enzyme genes as potential risk factors for AD, and being mindful of the limited statistical power of these analyses, we hypothesized that additional lysosomal enzyme genes will also represent risk factors for AD. Although deleterious mutations/variants in neither the β-glucuronidase (*GUSB*) nor the N-acetylglucosaminidase (*NAGLU*) genes appeared to be enriched in AD patients, we nonetheless determined the effects of heterozygosity of *GUSB* and *NAGLU* on ISF Aβ levels in C57Bl/6 mice and on Aβ plaques and insoluble Aβ in 5xFAD mice. Heterozygosity of *GUSB* decreases enzyme activity by approximately 50% (Fig. 5A). Similar to the *PPT1*, *IDUA*, and *GALC* genes, heterozygosity of *GUSB* has a gene-dosage effect on ISF Aβ levels in C57Bl/6 mice (Fig. 5B). In addition, heterozygosity of *GUSB* increased the plaque burden (Fig. 5C, Supplemental Fig. 5A) and levels of detergent-insoluble Aβ_40_ (Fig. 5D) and Aβ_42_ (Fig. 5E) in 5xFAD mice.

**Fig. 5.**
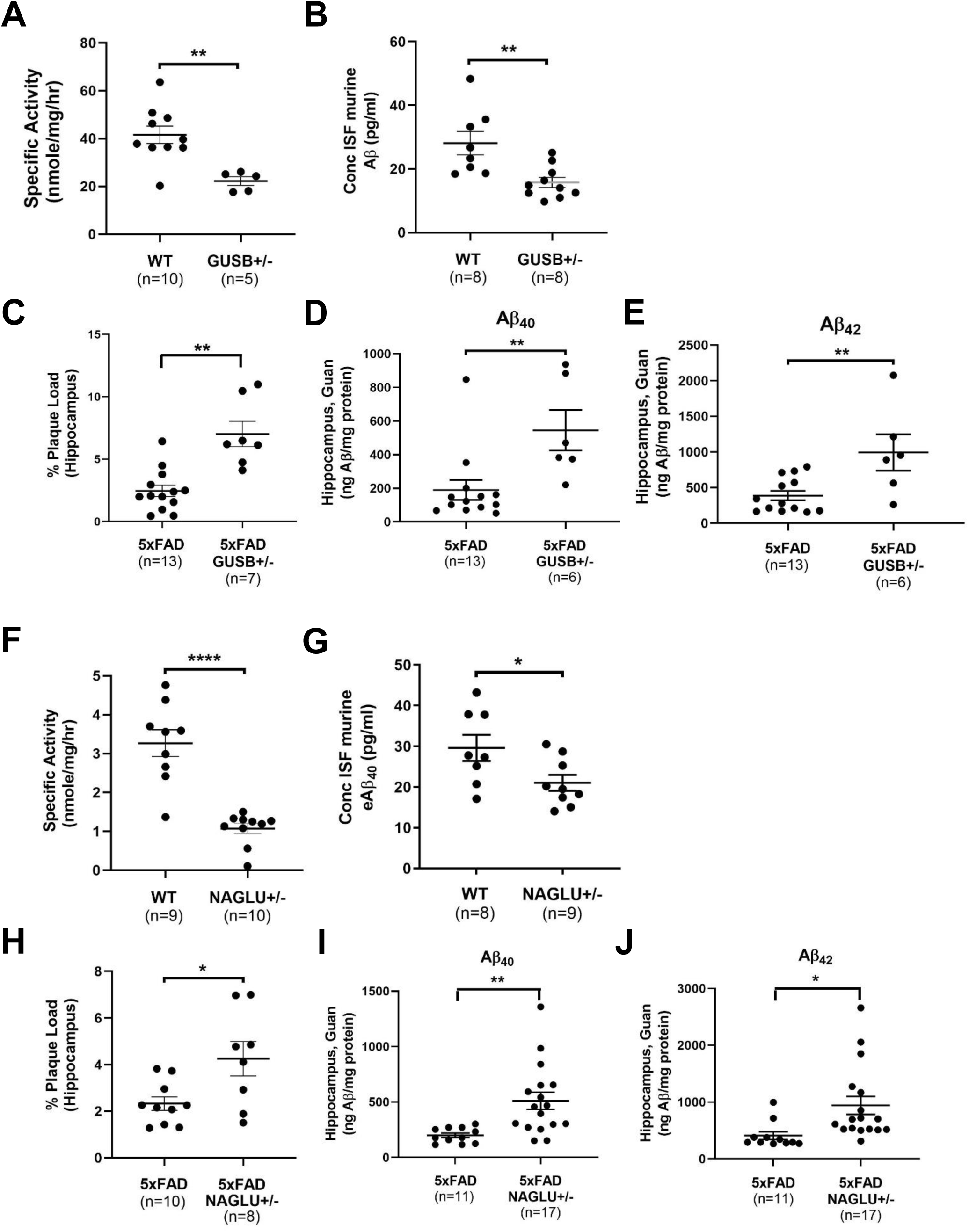
Two lysosomal enzyme genes that were <u>not</u> identified in the human genetic analysis affect Aβ metabolism and exacerbate the AD phenotype of 5xFAD mice. Three and one half-month-old mice that are heterozygous for the N-acetylglucosaminidase (*NAGLU*) gene (NAGLU+/-) have approximately half of the enzyme activity (A) and significantly decreased ISF Aβ levels (B) compared to age-matched wild type mice (WT). Heterozygosity of *NAGLU* also significantly increased the Aβ plaque load (C) and the level of detergent-insoluble Aβ_40_ (D) and Aβ_42_ (E) in 5xFAD mice. Three and one half-month-old mice that are heterozygous for the β-glucuronidase (*GUSB)* gene (GUSB+/-) have approximately 50% enzyme activity (F) and significantly decreased ISF Aβ levels (G) compared to age-matched wild type mice. Heterozygosity of *GUSB* also significantly increased the Aβ plaque load (H) and the level of detergent-insoluble Aβ_40_ (I) and Aβ_42_ (J) in 5xFAD mice. *p<0.05, **p<0.01, ****p<0.0001.

Heterozygosity of *NAGLU* decreased brain NAGLU enzyme activity (Fig. 5F) and has a gene-dosage effect on ISF Aβ levels in C57Bl/6 mice (Fig. 5G). Haploinsufficiency of NAGLU also significantly increased the plaque burden (Fig. 5H, Supplemental Fig. 5B) and levels of insoluble Aβ_40_ (Fig. 5I) and Aβ_42_ (Fig. 5J) in 5xFAD mice. Heterozygosity of *NAGLU* also significantly decreased the life span of 5xFAD mice (Supplemental Fig. 5C), although to a lesser extent than *PPT1*.

## Discussion

We show here that heterozygosity of known pathogenic mutations and predicted protein-damaging variants in at least five lysosomal enzyme genes are enriched in AD patients in case-controlled cohorts. Consistent with the hypothesis that lysosome dysfunction is associated with AD, human proteomic data identified the lysosomal storage disease pathway as being altered in AD patients. However, the proteomic data represent a snapshot in time and do not establish whether the changes in the lysosome pathway are causative or simply a response to disease. Therefore, three of the genes identified in the genetic analysis were validated *in vivo* by demonstrating that heterozygosity, by itself, affects Aβ trafficking in C57Bl/6 mice and exacerbates Aβ aggregation in a mouse model of AD. In fact, the number of lysosomal enzyme genes was expanded beyond the human genetic analysis by demonstrating that heterozygosity of *NAGLU* and *GUSB*, neither of which was identified in the genetic analysis, had similar effects on ISF Aβ levels and exacerbated the 5xFAD phenotype. That Aβ levels are impacted in lysosomal enzyme haploinsufficient mice expressing WT murine Aβ underscores that these changes are not related to using a transgenic mouse model harboring multiple human mutations. In total, at least seven different lysosomal enzyme genes have been identified that likely represent risk factors for AD when carrying heterozygous variants. Given that we validated two genes *in vivo* that were not identified in genetic analysis, the fact that the genetic analyses were underpowered relative to standard genetic linkage studies, and homozygous mutations in a number of other lysosomal enzyme genes affect Aβ metabolism, we predict that heterozygous deleterious mutations/variants in other lysosomal enzyme genes will also be associated with AD.

These findings are of particular interest since they challenge the dogma that, in general, carriers of lysosomal enzyme gene defects are normal throughout life. In fact, it is becoming increasingly apparent that this long-held belief may not be correct. Perhaps the best-known example of haploinsufficiency of a lysosomal enzyme gene being associated with an adult-onset neurological disease is the role of heterozygous missense variants in the *GBA* gene in PD (29). Carriers of coding variants in sphingomyelinase (*SMPD1*) are also associated with PD (39). There is also ample evidence from studies in experimental systems that heterozygous mutations in a number of lysosomal genes are associated with a Parkison-like phenotype (40). The association of heterozygous mutations in *GBA* is not the only example of carriers of lysosomal gene defects being associated with adult-onset neurological disease. Heterozygous mutations in *GALC*, *GRN*, and *NAGLU* are associated with multiple sclerosis, fronto-temporal dementia, and adult-onset axonal sensory neuropathy, respectively (30, 41, 42). Finally, a recent study in an extended family identified a severe heterozygous mutation in the *NPC1* gene that is associated with a dominant form of AD (31). These studies support the hypothesis that heterozygosity of deleterious variants in some, perhaps many, lysosomal genes predispose carriers to adult-onset neurodegenerative disorders.

At this point, these data do not allow us to determine if haploinsufficiency of a single lysosomal enzyme results in global lysosome dysfunction or in perturbations of a specific pathway affecting Aβ metabolism. However, the substrate specificity of some of the lysosomal enzymes might provide some insights. For example, PPT1 is responsible for removing the fatty acyl chains from palmitoylated proteins. Interestingly, several proteins involved in Aβ metabolism are palmitoylated proteins. The N-terminal region of APP is palmitoylated *in vitro* and *in vivo* and APP palmitoylation is involved in the amyloidogenic process (35). APP also interacts with the palmitoyl acyltransferase, DHHC12/AID, which suppresses APP trafficking by tethering APP in the Golgi and inhibiting APP metabolism (43). Palmitoylated APP is enriched in lipid rafts and is a more efficient substrate for BACE1. BACE1 is also located to lipid rafts based on it being palmitoylated, thus promoting Aβ production (44). The palmitoyltransferase inhibitors, 2-bromopalmitate and cerulenin, decrease Aβ production (45, 46). Inhibition of palmitoylated-APP formation by ACAT inhibitors (the preferred long-chain acyl substrate for ACAT1 is palmitoyl-CoA, the precursor of palmitate) would appear to be a valid strategy for prevention and/or treatment of AD (44). Therefore, a perturbation in the balance between palmitoylation and de-palmitoylation could affect Aβ metabolism and might be associated with AD. Although it is unclear exactly how *PPT1* heterozygosity affects the levels of the secretases, the fact that α-secretase (non-amyloidogenic) levels are depressed and β- and γ- secretase (amyloidogenic) levels are increased in aged heterozygous *PPT1* mice, and the AD pathway is activated in *PPT1* heterozygous sheep supports the role of PPT1 haploinsufficiency in AD.

Another example where perturbation of a specific degradative pathway might affect Aβ aggregation and/or trafficking is with the *NAGLU*, *IDUA*, and *GUSB* genes. These lysosomal enzymes are involved in the stepwise degradation of glycosaminoglycans. Specifically, NAGLU is required for the degradation of heparan sulfate (HS), IDUA is required for the degradation of HS and dermatan sulfate (DS), and GUSB is required for the degradation of HS, DS, and chondroitin sulfate (CS) (47). The common substrate for all three enzymes is HS. Therefore, all three enzymes are required to maintain homeostatic levels of heparan sulfated proteoglycans (HSPG). HSPGs regulate the oligomerization, clearance, endocytosis and trafficking of a variety of pathogenic proteins and have been shown to mediate cellular Aβ uptake and accelerate its oligomerization and aggregation (48, 49). HSPGs are also present in Aβ plaques (50, 51). These findings suggest that HS and HSPGs play important roles in Aβ metabolism and the pathogenesis of AD. Complete deficiency of NAGLU, IDUA, and GUSB result in the pediatric lysosomal storage diseases mucopolysaccharidosis IIIB (MPSIIIB), mucopolysaccharidosis I (MPSI), and mucopolysaccharidosis VII (MPSVII), respectively. Untreated MPS patients rarely live beyond their twenties and thus do not have AD. However, it has been shown that these patients have increased levels of soluble Aβ and accumulate intracellular Aβ in cells throughout the brains (21, 22). Data suggest that intraneuronal Aβ is neurotoxic and precedes extracellular deposition in both humans (52) and mouse models (53). In addition, pharmacological inhibition of HSPG binding of pathogenic proteins and genetic reduction of HSPG synthesis facilitate the clearance of pathogenic proteins and reduce their aggregation (54, 55). Therefore, it is possible that heterozygosity of *NAGLU*, *IDUA*, and *GUSB* might alter the normal levels of HSPGs and exacerbate Aβ aggregation. There are three additional lysosomal genes that are involved in heparan sulfate metabolism; heparan N-sulfatase (*SGSH*), α-glucosaminide acetyltransferase (*HGSNAT*), and N-acetylglucosamine 6-sulfatase (*GNS*). If haploinsufficiency of lysosomal enzymes responsible for proper HS metabolism result in subtle changes in the levels of HSPGs and are involved in the pathogenesis of AD, by extrapolation, heterozygous mutations in *SGSH*, *HGSNAT*, and *GNS* might also represent risk factors for AD.

It is less clear how heterozygosity of other lysosomal enzyme genes discovered in the human genetic analysis might be involved in AD. Galactocerebrosidase and acid ceramidase are lysosomal enzymes that catabolize glycolipids and were identified in the human genetic analysis. Homozygous deficiency of *GALC* and *ASAH1* result in the lipid storage diseases, globoid cell leukodystrophy (Krabbe disease) and Farber lipogranulomatosis, respectively. Interestingly, the substrate specificity of these two enzymes overlaps. *GALC* and *ASAH1* can both cleave galactosylceramide, albeit at different sites (56). Subtle perturbation in the levels or distribution of various galactosylceramide species could affect specific pathways involved in Aβ metabolism or disrupt the integrity of lipid membranes and thus result in secondary or global lysosome defects. In fact, it has been shown that accumulation of one of the substrates of GALC, galactosylsphingosine, can affect lipid microdomains and disrupt lipid membranes (57, 58).

With respect to the possibility that heterozygosity a single lysosomal enzyme might cause some sort of global lysosome dysfunction, it has been shown that complete deficiency of a single lysosomal enzyme can result in the accumulation of substrates other than those degraded by the deficient enzyme (59). Therefore, it is conceivable that heterozygosity of a single lysosomal enzyme could have effects on other enzymes and their substrates thus leading to a subtle but chronic global lysosome dysfunction. Another observation that would be consistent with global lysosome dysfunction is the fact that all five lysosomal genes validated *in vivo* significantly decreased the levels of Aβ in brain ISF. This common observation also appears to be counterintuitive since AD is characterized by the accumulation of extracellular Aβ plaques. However, as mentioned above, it has been shown in several distinct lysosomal storage diseases that there can be a massive accumulation of intracellular Aβ in the absence of extracellular plaques. This might suggest that a severe (in the case of homozygous deficiency) or a subtle (in the case of haploinsufficiency) perturbation in lysosome function affects some common pathway resulting in the sequestration of Aβ within the cell. This would also be consistent with a defect in the normal trafficking or secretion of Aβ fragments as described above.

Finally, the fact that soluble lysosomal enzymes can be secreted from a cell and taken up by adjacent cells through a well-established, receptor-mediated mechanism, commonly referred to as cross-correction (60), opens the possibility that lysosomal enzyme-associated forms of AD might be amenable to treatment. The biochemical, histological and clinical improvements observed in the 5xFAD/PPT1+/- mice following CNS-directed, AAV-mediated gene therapy strongly support this contention. Complementing the haploinsufficient enzyme could be accomplished by a number of means, including direct gene therapy, *ex vivo* stem cell-directed gene therapy, protein replacement therapy, chaperone therapy, etc. Several of these approaches are already FDA-approved or are currently in clinical trials for homozygous lysosomal enzyme deficiencies.

Cumulatively, these data strongly suggest that heterozygous deleterious mutations/variants in lysosomal enzyme genes are associated with AD. However, at this point it is difficult to determine the exact mechanism/s involved or the overall risk. In this study, we identified at least seven different genes that might represent risk factors for AD and we believe that this underestimates the total number that are associated with AD. In addition, there are 40 or more LSD that have a significant neurological component. Even if the risk associated with an individual lysosomal enzyme gene is low, the number of genes involved could be relatively large. For example, based on an overall incidence of LSD of 1/7000 - 1/5000 live births (61), the carrier frequency for a lysosomal gene defect in the general population would be 1/41 - 1/35. For an individual LSD with an incidence of 1/100,000 live births, the carrier frequency would be 1/158. Therefore, heterozygous deleterious mutations/variants in lysosomal enzyme genes could be associated with a significant proportion of AD patients.

## Materials and methods

### Animals and human samples

All the mice used in this study were maintained on a congenic C57BL/6J background by MSS at Washington University School of Medicine. The PPT1-deficient mouse model used for this study was first generated by Gupta et al (34), and is a well-established model of CLN1 disease, recapitulating most of the human behavioral and pathological phenotypes of the human disease. The *NAGLU*-deficient mouse model (MPSIIIB) was generated by Li et al (62). The *GALC*- and *GUSB*-deficient mice were spontaneously arising mutants that have been thoroughly characterized (63, 64). The mouse model of mucopolysaccharidosis type I was created by targeted disruption of the α-L-idurondase gene (65). The 5XFAD (line Tg6799) mouse model of AD (36) was purchased from the Jackson Laboratory. The colonies were maintained separately through sibling mating. Both male and female mice were used in this study. All procedures were performed in accordance with NIH guidelines as well as a protocol approved by the Institutional Animal Care and Use Committee (IACUC) at Washington University School of Medicine.

The PPT1 sheep were maintained at the Roslin Institute (Edinburgh, Scotland) with free access to food and water. The *PPT1* heterozygous and normal control animals were placed under surgical anesthesia and perfused with normal saline at the ages indicated (Supplemental Table 3). The brains were removed and flash-frozen in liquid nitrogen. All the work involving sheep was reviewed and approved by the Animal Welfare and Ethical Review Board (AWERB) at the Roslin Institute and conducted under the authority of the UK Home Office. Use of human tissue for post-mortem studies has been reviewed and approved by the Edinburgh Brain Bank Ethics committee and the ACCORD Medical Research Ethics Committee. The Edinburgh Brain Bank is a Medical Research Council funded facility with Research Ethics Committee (REC) approval.

### Human Genetic Analysis

The participants with WGS data in the ADSP R4 release, known as the R4 ∼36k WGS Project Level VCF, were the focus of the current analysis. All variants were excluded based on the low complexity region (LCR), monomorphism, ABHet ratio (ABHet < 0.25 or ABHet > 0.75), and variant with high read depth (>500 reads). SNPs and INDELs were filtered out based on various hard filters criteria: (1) quality depth (QD ≥ 7 for indels and QD ≥ 2 for SNPs), (2) Mapping quality (MQ ≥ 40), (3) FisherStrand (FS) balance (FS ≥ 200 for indels and FS ≥ 60 for SNPs), (4) Strand Odds Ratio (SOR ≥ 10 for Indels and SOR ≥ 3 for SNPs), (5) Inbreeding Coefficient (IC ≥ −0.8 for indels), and (6) ReadPosRankSum Test for relative positioning of reference vs. alternative alleles within reads (RPRS ≥ −20 for Indels and RPRS ≥ −8 for SNPs). Only variants that had a filter “PASS” in the vcf file were included in the analysis.

Plink (Plink v1.9) software was used to perform the quality check at the variant level and subject level. In case-control datasets, variants were removed based on missingness rate higher than 2% and individuals were removed based on missingness higher than 2%. Variants that had a Hardy Weinberg equilibrium (p < 1 × 10^−30^) were excluded. Individual relatedness samples were filtered out based on the given set of ADSP R4 related samples list. Principal Component Analysis (PCA) was calculated using pruned variants together with the population data from 1000 genomes and the analysis was restricted to only samples from Central European and African American ancestry.

Functional annotation was performed using the dbSNP, dbNFSP and with minor allele frequencies from gnomAD, ExAC and 1000 genomes. All the variants were annotated using SnpEff and SnpSift. Protein-coding variants were annotated as missense variant, protein-altering variant, frameshift variant, inframe deletion, inframe insertion, splice donor variant, splice acceptor variant, start lost, stop gained, and stop lost. Pathogenic and likely pathogenic variants for selected endolysosomal genes were retrieved from ClinVar database (https://ftp.ncbi.nlm.nih.gov/pub/clinvar/vcf_GRCh38/archive_2.0/2023/). For gene-based analysis, the exonic variants having CADD score (32) greater than 30 and all selected ClinVar pathogenic variants were included. The SKAT package was used to perform SKAT-O and Burden tests using dichotomize traits adjusted for sex and PCs (PC1-PC10) for gene based analysis. In addition, pathway level analysis was performed on the ClinVar pathogenic variants using the SKAT-O test adjusted for sex and PCs (PC1-PC10).

### Protein extraction and proteomic analyses

Tissue samples from humans and sheep (Supplemental Tables 2 and 3) were used for proteomic analysis as follows. The brain tissue slices were resuspended in an extraction buffer containing 5% SDS, 50mM Triethyl Ammonium Bicarbonate (TEAB), pH 8.5 at sample to buffer ratio of 1:10 (w/v) and homogenized using Precellys homogenizer at 5,000g for 20 seconds in a ceramic beads vial (Precellys Lysing Kit, Tissue homogenizing CK mix). The extracts were centrifuged for 10 min at 16,000g and supernatant was sonicated for 10 cycles with 30 sec on and 30 sec off per cycle on a Bioruptor Pico Sonicator (Diagenode). After sonication, samples were centrifuged (16,000xg for 10min), supernatant collected, and a BCA assay was performed.

The protein samples were digested on the S-Trap micro column (Protifi, USA) following manufacturer’s protocol with minor modifications. Briefly, 20 μg protein in extraction buffer was reduced and alkylated using 10 mM dithiothreitol and 40 mM iodoacetamide respectively, at 45°C for 15 min. Alkylation was stopped by adding phosphoric acid to a final concentration of 2.5%. The protein solution was diluted six-fold with binding buffer (90% v/v, Methanol in 100 mM TEAB), vortexed gently, and loaded into an S-Trap micro column. After centrifugation at 4,000 g for 1 min, the column was washed three times with 150 μL binding buffer. Proteolytic digestion was carried out by adding 20 μL of digestion buffer containing 1μg trypsin in 50 mM TEAB. The column was incubated at 47°C for 2 hours. Digested peptides were eluted using 40 μL of three buffers consecutively: (i) 50 mM TEAB, (ii) 0.2% (v/v) Formic acid in H_2_O, and (iii) 50% (v/v) acetonitrile. Eluted peptides were pooled and cleaned up using C_18_ stage tips and dried under a vacuum.

Purified peptides were separated over a 70 minute gradient on an Aurora-25 cm column (IonOpticks Australia) using a UltiMate RSLCnano LC System (ThermoFisher Scientific) coupled to a timsTOF FleX mass spectrometer via a Captivespray ionization source. The gradient was delivered at a flow rate of 200 nL/min and the column temperature was set at 50°C. Data were acquired using a diaPASEF method consisting of 12 cycles including a total of 32 mass width windows (27.2 Da width, from 400 to 1201 Da) with 2 mobility windows each, covering a total of 64 windows (1/K_0_ from 0.75 to 1.28 V s/cm^2^). Optimal m/z and ion mobility windows were designed with py_diAID utility using a DDA-PASEF run previously acquired from a pool of the analyzed samples. The TIMS elution voltage was linearly calibrated to obtain 1/K_0_ ratios using three ions from the ESI-L Tuning Mix (Agilent, Santa Clara, CA, USA) (*m/z* 622, 922, 1222) using timsControl (Bruker). For DDA-PASEF acquisition, the full scans were recorded from 100 to 1700 m/z spanning from 1.45 to 0.65 Vs/cm2 in the mobility (1/K0) dimension. Up to 10 PASEF MS/MS frames were performed on ion-mobility separated precursors, excluding singly charged ions which are fully segregated in the mobility dimension, with a threshold and target intensity of 1750 and 14,500 counts, respectively. In both DDA- and DIA-PASEF modes, the collision energy was ramped linearly from 59 eV at 1/k0 = 1.6 to 20 eV at 1/k0 = 0.6.

Raw mass spectral data were processed using Spectronaut Version 18.0 (Biognosys, Schlieren, Switzerland) with a direct DIA method. As a first step, database search was performed by pulsar search engine against either Uniprot Human (20,601 entries) or Sheep (21,220 entries) sequence database. The search parameters included the use of trypsin/P as the enzyme with a specific digestion type, a 7–52 peptide length range and up to two missed cleavages. Oxidation of methionine and acetylation of protein N-terminal were variable modifications and carbamidomethyl of cysteines was a fixed modification. A false discovery rate (FDR) of 1% was applied at peptide and protein group identifications. The generated spectral library was then used by Spectronaut for DIA analysis and quantity was determined at MS2 level using the area of extracted chromatogram traces. An automatic local cross-run normalization strategy was followed, and the MaxLFQ method was used for protein quantification.

Seven thousand nine hundred and fifteen proteins were identified in the AD patients and 8835 in the PPT1 ^+/-^ sheep, respectively. AD patient samples were run in triplicate and only proteins identified in 2 of the 3 replicates and more than 1 unique peptide were included in the analysis moving forwards. Imputation of missing values was not implemented. Samples from *PPT1* heterozygous sheep with 3 or more replicates that were identified by more than 1 unique peptide were included in the analysis moving forwards. The median of each group control versus AD or Het was calculated. Thereafter, the ratio of AD or PPT1+/- divided by the control was calculated to determine protein abundance as a ratio relative to wild type. In the context of Het sheep, PPT1 protein was used as an internal control where the median of PPT1^+/-^ samples calculated as 35% decrease from WT controls.

### Proteomic bioinformatics analysis

Given the nature of the questions investigated, not knowing what a meaningful threshold for magnitude of change should be for protein alterations within a given defined molecular pathway/cascade data was analysed and differing abundance thresholds are indicated accordingly (Table 2). Data were further analyzed with Ingenuity Pathway Analysis (IPA) software (Ingenuity Systems, Silicon Valley, CA, USA) to explore the cellular and molecular pathways that may have been altered as a consequence of differential expression in Alzheimer’s Disease (AD) human post mortem samples or heterozygous PPT1 +/- sheep (Het) versus their respective controls. A right-tailed Fisher’s Exact Test was used to calculate the *p*-value determining the probability that each cellular and molecular function or canonical pathway assigned to that dataset is due to chance alone, and the final lists of functions and pathways were ranked accordingly to the resulting *p*-value.

The same datasets of proteins were used for network generation in IPA. Each identifier was mapped to its corresponding entry in Ingenuity’s Knowledge Base, and these proteins were overlaid onto a global molecular network developed from information contained in the Ingenuity Knowledge Base as previously described (66). Networks were then algorithmically generated based on their connectivity. The Functional Analysis of each network identified the biological functions and/or diseases that were most significant to the proteins in the network. A right-tailed Fisher’s Exact Test was used to calculate the *p*-value representing the probability that the overlaps reported between the input dataset and the base pathway/network (e.g., all of the constituent components of RhoGD signalling) are due to chance alone. The resulting networks are a graphical representation of the molecular relationships between proteins, where proteins are represented as nodes, and the biological relationship between two nodes is represented as an edge (line). All edges are supported by at least one reference from the literature, from a textbook or from canonical information stored in the Ingenuity Knowledge Base, all supporting an *in vivo* or *in vitro* observation of a protein–protein or protein–DNA interaction (as opposed to a merely predicted interaction from in silico experimentation).

### AAV9-hPPT1 vector and intracranial injections

The AAV9-hPPT1 vector used in this study was initially created and characterized by Griffey et al. (2004) (67). The concentration of the AAV vector was 1 x 10^12^ vg/ml in sterile lactated Ringer’s solution. For the intracranial injections in 3.5-month-old mice, animals were placed under surgical anesthesia with isofluorane, the head shaved, and cleaned with 70% isopropanol and Betadine. The animals were placed in a three-point stereotactic frame and a midline incision was made under aseptic conditions. A 2mm burr hole was made and the AAV vector slowly injected. Four, 2μl intraparenchymal bilateral injections were performed at the following coordinates from the bregma to target the cortex (−1.0mm – AP, −2.5mm – ML, −0.7 - DV) and hippocampus (− 2.0mm - AP, −1.5mm – ML, −1.5mm – DV). The single intracerebroventricular injections contained the identical dose (8 x 10^9^ vg) of virus and were injected at the following coordinates: left hemisphere, −0.5mm – AP, −1.0mm – ML, −1.7mm – DV. Once the injection was complete, the needle was removed and the incision closed with 5- 0 nylon suture. The animals were allowed to recover on a temperature-regulated pad and returned to their home cage. AAV injections in newborn animals were performed identically as described previously (66).

### Behavioral analyses

The spontaneous alternation Y-maze and elevated plus maze were performed according to previously published procedures (68). For the Y-maze, a mouse was placed in the center of a Y-maze that contained three arms that were 10.5 cm wide, 40 cm long and 20.5 cm deep oriented at 120° with respect to each successive other arm. Mice were allowed to explore the maze for 8 min and entry into an arm was scored only when the hindlimbs had completely entered the arm. An alternation was defined as any three consecutive choices of three different arms without re-exploration of a previously visited arm. Dependent variables included the number of alternations and arm entries along with the percentage of alternations, which was determined by dividing the total number of alternations by the total number of entries minus 2, then multiplying by 100.

In the elevated plus maze, a 5 min trial was conducted in a dark room using a standard mouse elevated plus maze from Kinder Scientific (Poway, CA). The maze consisted of a black acrylic surface measuring 5×5cm and elevated 63cm above the floor equipped with photo beam pairs to track the position of the mouse. Four arms (35cm long, 5cm wide; two open and two with 15cm high walls) extended from a central area. The MotorMonitor software (Kinder Scientific, LLC, Poway, CA) quantified beam breaks as duration, distance traveled, entries, and time at rest in each zone (closed arms, open arms and center area). Variables analyzed were percent of total time, distance, and entries into the open arms of the maze during the trial, as well as total distance traveled.

### *In vivo* Aβ microdialysis

To assess ISF Aβ levels in the hippocampus of awake, freely moving WT, *PPT1*+/-, *GALC*+/-, *NAGLU*+/-, *IDUA*+/-, and *GUSB*+/- mice over time *in vivo*, microdialysis was performed as previously described (69). Briefly, under isoflurane anesthesia, a guide cannula was stereotaxically implanted above the hippocampus (3.1 mm behind bregma, 2.5 mm lateral to midline, and 1.2 mm below dura at a 12° angle). A microdialysis probe was inserted through the guide cannula into the brain. Artificial CSF (1.3 mM CaCl_2_, 1.2 mM MgSO_4_, 3 mM KCl, 0.4 mM KH_2_PO_4_, 25 mM NaHCO_3_, and 122 mM NaCl, pH 7.35) containing 0.15% bovine serum albumin (BSA) (Sigma) filtered through a 0.22 μm membrane was used as perfusion buffer. Flow rate was a constant 1.0 μl/min. Samples were collected every 60 to 90 min overnight, which gets through the 4- to 6-hour recovery period, and the mean concentration of Aβ over the 6-hour preceding treatment was defined as basal concentration of ISF Aβ. Samples were collected through a refrigerated fraction collector and assessed for Aβ40 or Aβ42 by ELISAs [β-amyloid antibodies mHJ5.1 and mHJ2 (kind gift from David Holtzman, Washington University School of Medicine) were used for the ELISA assays]. The mean concentration of exchangeable Aβ (eAβ) detected was corrected for assuming a 12% recovery rate of Aβ into the microdialysis probe (37). The absolute concentration of ISF Aβ for each animal was the mean concentration of six hours of collection between hours 10-16 after probe insertion. Analysis of microdialysis experiments used unpaired t-tests to compare two genotypes or a one-way ANOVA with an appropriate correction for multiple comparisons to compare three genotypes.

### Immunohistochemistry and plaque quantification

Immunohistochemstry and subsequent quantification were performed essentially as described (69). Mice were anesthetized with an intraperitoneal injection of pentobarbital, followed by transcardially perfusion with ice-cold phosphate buffer saline (PBS) with 0.3% heparin. The brains were carefully removed and separated into two hemispheres along the mid-sagittal plane. The left hemisphere was fixed in 4% paraformaldehyde in phosphate buffer overnight and then transferred to 30% sucrose in PBS and allowed to sink at 4°C before sectioning. The right hemisphere was dissected into regions and frozen at −80°C until use. Coronal brain sections 50 μm wide were sliced in 300 μm intervals using a freezing sliding microtome.

Brain sections (50 μm) were collected every 300 μm from rostral anterior commissure to caudal hippocampus. Sections were immunostained with biotinylated mHJ3.4 antibody (kind gift from David Holtzman, Washington University School of Medicine). Four slices per animal were used corresponding to Paxinos and Franklin Mouse Brain Atlas, Second Edition, bregma −1.7, −1.94, −2.3, and −2.54. NanoZoomer Digital Scanner (Hamamatsu Photonics, Bridgewater, NJ) was used to create high-resolution digital images of the stained brain slices. The total area of plaque coverage in the hippocampus was measured using ImageJ software (National Institutes of Health) and expressed as percentage total area for each slice.

### Western Blotting

Immunoblotting (Western) was performed essentially as described with minor changes (70). Brain tissue (mouse hippocampus) was homogenized in ice-cold lysis buffer (50 mM Tris-HCl, 150 mM NaCl, 1% Nonidet P- 40, 0.5% Triton X-100) supplemented with protease and phosphatase inhibitors (A32965 and A32957, Thermo Scientific, Waltham, MA, USA), incubated on ice for 20 min, and centrifuged at 16,000 x g for 20 min at 4°C. Protein concentration was determined using Pierce BCA Protein Assay Kit (23227, Thermo Scientific) according to manufacturer’s instructions. 20 ug of protein were loaded per lane and separated on SDS-PAGE (12% (#3450117) and 4-12% (#3450123) Criterion XT Bis-Tris gels, Bio Rad, Hercules, CA, USA), and transferred onto PVDF membrane (1620177, Bio Rad) using semi-dry Trans-Blot® Turbo™ protein transfer system (1704150, Bio Rad) followed by blocking for 1 h at RT in 5% skimmed milk in Tris-buffered saline with 0.05% Tween-20 (P7949, MilliporeSigma, St. Louis, MO, USA) (TBST). All primary antibody incubations were performed on a shaker overnight at 4°C. The following primary antibodies and dilutions were used: anti-ADAM10 [B-3] (1:1000, sc-28358, Santa Cruz Biotechnology, Dallas, TX, USA), β-amyloid precursor protein [CT695] (1:1000, 51-2700, Thermo Scientific), anti-BACE1 [D10E5] (1:500, 5606, Cell Signaling Technology, Danvers, MA, USA), Presenilin-1 [CT, clone PS1-loop] (1:1000, MAB5232, MilliporeSigma) and anti-GAPDH (1:4000, MA5-15738, Thermo Scientific). After removing the primary antibodies, the membranes were washed 3 x 5 min with TBST at RT followed by secondary antibody incubation. Horseradish peroxidase-conjugated anti-mouse, anti-rabbit IgG (7076S and 7074S, Cell Signaling Technology) or anti-goat IgG (A27014, Thermo Scientific) secondary antibodies were diluted 1:5000 in 5% skimmed milk in TBST and incubated for one hour at room temperature. Signals were visualized using Clarity Western ECL Substrate (#1705061, Bio Rad). The blots were imaged using ChemiDoc^TM^ Imaging System (#12003153, Bio Rad). Band intensity was quantified using ImageJ software (National Institutes of Health) and normalized to GAPDH signal, which was used as loading control. The values shown are the normalized band intensities relative to the experimental control group. Mean ± SEM per group are shown.

### Lysosome Enzyme Assays

The lysosomal enzyme assays were performed essentially as described previously (67). Experimental animals and controls were sacrificed via lethal injection (Fatal-Plus; Vortech Pharmaceuticals) at the times identified in the figure legends. The brains were removed and bisected sagittally. The striatum was carefully dissected and flash frozen. The tissue samples were homogenized in buffer containing 10 mM Tris (pH 7.5), 150 mM NaCl, 1 mM dithiothreitol, and 0.2% Triton X-100 and centrifuged at 20,000 rpms for 1 min at 4°C. Following centrifugation, the supernatant was removed and used for PPT1, GUSB, IDUA, NAGLU, and GALC enzyme assays using the respective 4-Methylumbelliferyl (4MU)-conjugated fluorometric substrates [PPT1 – 4MU-6-thio-palmitate-β-D-glucopyranoside (Cayman Chemical Co, #19524), GALC – 6HMU-β-D-galactoside (Carbosynth, #EH05989), NAGLU – 4MU-N acetyl-β-D-glucosaminide (Sigma, #M2133), GUSB – 4MU-β-D- glucuronide (Sigma, #M9130), IDUA – 4MU-α-L-iduronide (Sigma, #9531)]. Fluorescence was measured at 360nm excitation and 440nm emission wavelengths. Protein concentrations were determined by the coomasie dye-binding assay (BioRad) and enzyme activity was normalized to total protein.

### ELISA quantification

Aβ species in ISF and contralateral hemisphere were fractionated into 1% Triton X-100 soluble and insoluble (5M guanidine) fractions and detected and quantified by sandwich ELISA as previously published (69). Briefly, Aβx-40 and Aβx-42 peptides were captured with mouse monoclonal-coating antibodies HJ2 (anti-Aβ35-40) and HJ7.4 (anti-Aβ37-42). HJ5.1 (anti-Aβ13-28), a biotinylated antibody targeting the central domain, was used as the detecting antibody, followed by streptavidin-poly-HRP-40. Recombinant human Aβ1-40 or Aβ1-42 peptide was used to generate the standard curves for each assay. The ELISA assays were developed using Super Slow ELISA TMB (Sigma) with absorbance read on a BioTek Epoch plate reader at 650 nm. The mean ± SEM per group is shown.

### Statistical analyses

Two-tailed unpaired *t*-tests were used to compare between two groups. One-way or two-way ANOVA was used when comparing one or two independent variables, respectively, between multiple groups. The appropriate correction for multiple comparisons was used (Sidak, Tukey, or Bonferroni). Values were accepted as significant if *p* ≤ 0.05. Data in figures are presented as mean ± SEM. Prism 8 (GraphPad, San Diego, CA) was used for all statistical analyses. All measurements were performed blind to genotype.

## Acknowledgments

We would like to thank Marie Nunez and Kevin O’Dell for expert technical assistance. This work was supported in part by NIH grant NS100779, a grant from the Knight ADRC (P30AG066444), and a research contract from Jaya Biosciences to MSS. It was also supported by NIH grant RF1 AG079867 and R01 AG064902 to JRC. This work was supported in part by a grant from the McDonnell Foundation, pilot funding from the Hope Center for Neurological Disorders, the Danforth Foundation Challenge, and the Skandalaris Center at Washington University to BAB and MSS. TMW, SLE, and DTK acknowledge strategic investment from the Biotechnology & Biological Sciences Research Council, including Institute Strategic Program Grant funding (BB/X010945/1) and NIH R01 NS124655 to Jonathan Cooper (Washington University) for access to banked sheep tissue. We thank CS for access to the Human Tissue Bank (Affiliation) for the AD and human control samples and the Wellcome Trust for supporting the AD proteomic data generation through the One Health PhD Scheme to SLE. We thank the Genome Technology Access Center in the Department of Genetics at Washington University School of Medicine for help with genomic analysis. The Center is partially supported by NCI Cancer Center Support Grant #P30CA91842 to the Siteman Cancer Center and by ICTS/CTSA Grant# UL1TR002345 from the National Center for Research Resources (NCRR), a component of the National Institutes of Health (NIH), and NIH Roadmap for Medical Research. The intracranial injections were performed at the Hope Center Animal Surgery Core at Washington University School of Medicine. Behavior testing was performed by the Washington University Animal Behavior Core which is supported by funds provided by the McDonnell Center for Systems Neuroscience, McDonnell Center for Cellular and Molecular Neuroscience, and the Taylor Family Institute at Washington University in St. Louis. This publication is solely the responsibility of the authors and does not necessarily represent the official view of NCRR or NIH.

