## Supplemental Data for "Haploinsufficiency of lysosomal enzyme genes in Alzheimer’s disease"

**Supplemental Table 1** Pathogenic mutations and protein-damaging variants in the *GAA*, *IDUA*, *PPT1*, *GALC*, and *ASAH1* genes are enriched in AD patients of Central European (CEU) descent and *IDUA* and *GALC* are enriched in AA patients. The genomic position, reference and alternate nucleotides, rsIDs, annotation and CADD scores for each mutation/variant are indicated. The pathogenic mutations were identified in the ClinVar data base and the protein-damaging variants have a CADD score greater than 30. *Indicates variants that are identified as pathogenic in ClinVar but have a CADD score less than 30.

| Gene | Chromosome | Pos | Ref | Alt | rsIDs | Annotation | CADD Score |
|---|---|---|---|---|---|---|---|
| GAA | 17 | 80104542 | T | G | rs386834236 | c.-32-13T>G | *5.74 |
| | 17 | 80104837 | T | TC | rs761317813 | p.Asn87fs | *22.6 |
| | 17 | 80105110 | CT | C | rs386834235 | p.Glu176fs | *0.364 |
| | 17 | 80105857 | G | A | rs370950728 | p.Gly219Arg | *28.2 |
| | 17 | 80107818 | G | A | rs121907945 | p.Gly293Arg | 32 |
| | 17 | 80108495 | C | T | rs755253527 | p.Pro361Leu | *25.9 |
| | 17 | 80110837 | G | A | rs140826989 | p.Trp516* | 46 |
| | 17 | 80110841 | G | C | rs770780848 | c.1551+1G>C | 33 |
| | 17 | 80112081 | G | A | rs991082382 | p.Glu579Lys | *26 |
| | 17 | 80117110 | G | C | rs780655708 | c.2331+1G>C | 35 |
| IDUA | 4 | 1000880 | A | G | rs777295041 | c.386-2A>G | *28.4 |
| | 4 | 1002474 | T | C | | p.Leu393Pro | 32 |
| | 4 | 1003418 | C | G | rs121965021 | p.Pro533Arg | *24.6 |
| | 4 | 1003607 | A | G | rs753905054 | p.Asp570Gly | 33 |
| | 4 | 1004011 | G | C | rs1249951282 | c.1728-1G>C | 33 |
| | 4 | 1004329 | C | T | rs886043347 | p.Ser633Leu | *25 |
| | 4 | 987858 | C | T | rs121965020 | p.Gln70* | 39 |
| PPT1 | 1 | 40076841 | C | A | rs878853929 | c.798+1G>T | 35 |
| | 1 | 40076915 | T | A | rs386833664 | c.727-2A>T | 35 |
| | 1 | 40091398 | T | A | rs137852695 | p.Arg122Trp | *27.4 |
| | 1 | 40092397 | C | T | rs796052923 | c.234+1G>A | 33 |
| | 1 | 40097210 | A | T | rs137852699 | p.Leu10* | 36 |
| GALC | 14 | 87941559 | C | T | rs1182103005 | c.1671-1G>A | 33 |
| | 14 | 87950724 | G | A | rs770485731 | p.Arg396Trp | *25.3 |
| | 14 | 87963474 | G | C | | p.Tyr357* | 38 |
| | 14 | 87965504 | C | T | | c.1033+1G>A | 33 |
| | 14 | 87965534 | T | C | rs757407613 | p.Tyr335Cys | *28.7 |
| | 14 | 87968375 | G | A | rs780750448 | p.Arg290Cys | 32 |
| | 14 | 87986604 | T | C | | c.329-2A>G | 34 |
| | 14 | 87988206 | G | A | rs201422931 | p.Pro89Leu | 32 |
| | 14 | 87988484 | G | A | rs73312829 | p.Arg79Cys | 31 |
| | 14 | 87993029 | C | A | rs751975987 | p.Asp46Tyr | 31 |
| ASAH1 | 8 | 18059451 | G | A | | p.Gln311Ter | 36 |
| | 8 | 18061738 | TC | T | rs1369327940 | p.Gly233fs | *- |
| | 8 | 18083995 | G | A | rs756041561 | p.Gln22* | *18.57 |

**Supplemental Table 2**

Samples from Alzheimer's patients (AD) and controls (Ctrl) ranging from 72-93 years-of-age were collected from Superior temporal gyrus (BA41/42) brain region.

| Sample ID | Gender | Age (years) | Braak score | Patient status |
|---|---|---|---|---|
| SD007/23 | Female | 93 | 6 | AD |
| SD024/22 | Male | 75 | 6 | AD |
| SD040/21 | Female | 72 | 6 | AD |
| SD033/22 | Male | 93 | N/A | Ctrl |
| SD042/18 | Female | 73 | N/A | Ctrl |
| SD024/17 | Male | 72 | N/A | Ctrl |

**Supplemental Table 3**

*PPT1* heterozygous (Het) and wild type control (WT) samples were obtained from the entorhinal cortex brain region from the sheep ranging from 500-2022 days-of-age. Samples were obtained under Home Office license PP2318334 and taken from the Roslin Institute Tissue Bank.

(n=5 control and 4 Het PPT1 +/- sheep)

| Animal ID | Sex | Genotype | Age (days) |
|---|---|---|---|
| 25683 | F | WT | 1335 |
| 16A126 | M | WT | 500 |
| 16A174 | F | WT | 572 |
| 25685 | F | WT | 1335 |
| 25622 | F | WT | 1335 |
| 18A084 | F | HET | 1682 |
| 18A095 | F | HET | 554 |
| 17A018 | F | HET | 881 |
| 17A019 | M | HET | 2022 |

**Supplemental Figure Legends**

**Supplemental Fig 1:**

Representative micrographs of the frontal cortex from one of the AD patients listed in Supplemental Table 2. A) Immunohistochemistry shows extensive Ab plaques throughout the frontal cortex (Biolegend, anti-Ab antibody, 4G8 clone, 1:16,000 dilution). B) There is also extensive Tau staining in the same region of the frontal cortex (ThermoFisher, anti-Tau antibody, AT8 clone, 1:1000 dilution). Scale bar = 100mm.

**Supplemental Fig 2:**

Proteomic changes in the brains of AD patients converge on the lysosomal storage disease pathway. The levels of 30 lysosomal proteins are changed in the brains of AD patients in a similar pattern to that seen in lysosomal storage diseases. The colors and format of the symbols and lines are described in the accompanying legend. The shapes of the various symbols can be found here: https://qiagen.my.salesforce-sites.com/KnowledgeBase/articles/Knowledge/Legend.

**Supplemental Fig 3:**

Proteomic changes in the brains of heterozygous *PPT1* sheep are consistent with changes commonly observed in AD patients. The colors of the symbols and lines, as well as the line styles are described in the accompanying legend. The shapes of the various symbols can be found here: https://qiagen.my.salesforce-sites.com/KnowledgeBase/articles/Knowledge/Legend.

**Supplemental Fig 4:**

The elevated plus maze behavioral test was performed on 5xFAD/PPT1+/- mice treated at 3.5 months with intracerebroventricular AAV9-GFP (GFP ICV), intracerebroventricular AAV9-hPPT1 (PPT ICV), or intraparenchymal (PPT IP). Although there was an apparent improvement in the mean performance of the PPT ICV and PPT IP animals compared to the GFP ICV animals, the differences were not statistically significant (ns).

**Supplemental Fig 5:**

There is an obvious increase in the number and distribution of Ab plaques in seven-month-old 5xFAD/IDUA+/-, 5xFAD/GALC+/-, and 5xFAD/GUSB+/- animals compared to age-matched 5xFAD mice (A) (Note: The Ab immunostaining for the 5xFAD, 5xFAD/IDUA+/-, 5xFAD/GALC+/-, and 5xFAD/GUSB+/- animals was performed at the same time with the same reagents to ensure consistency for direct comparisons). Those increases are statistically significant (see Fig 4C, 4H and 5A, respectively). There is an obvious increase in the number and distribution of Ab plaques (B) in seven-month-old 5xFAD/NAGLU+/- mice compared to age-matched 5xFAD mice (B). That increase is statistically significant (see Fig 5H). There is also a significant decrease in life span compared to either 5xFAD or NAGLU+/- mice (C).

**Supplemental Figure 1**

A

B

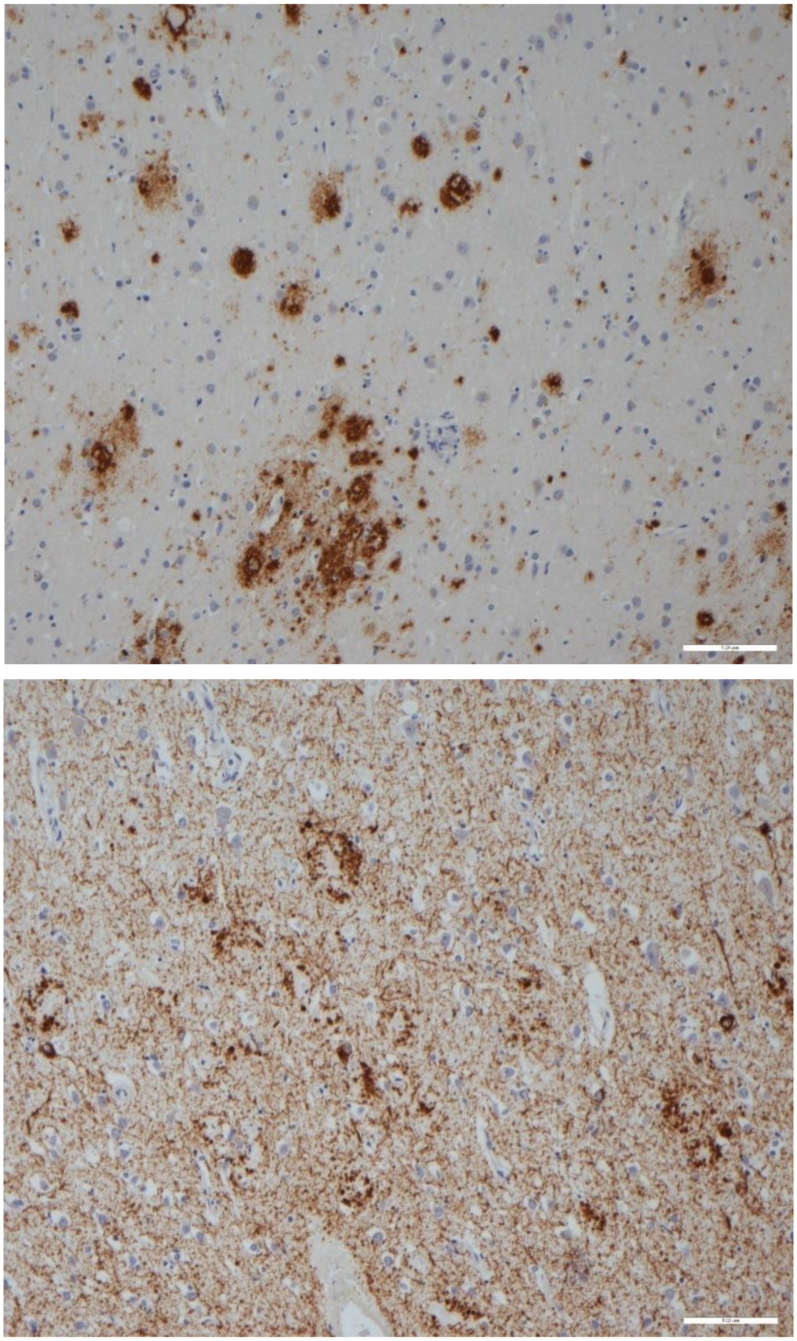

**Supplemental Figure 2**

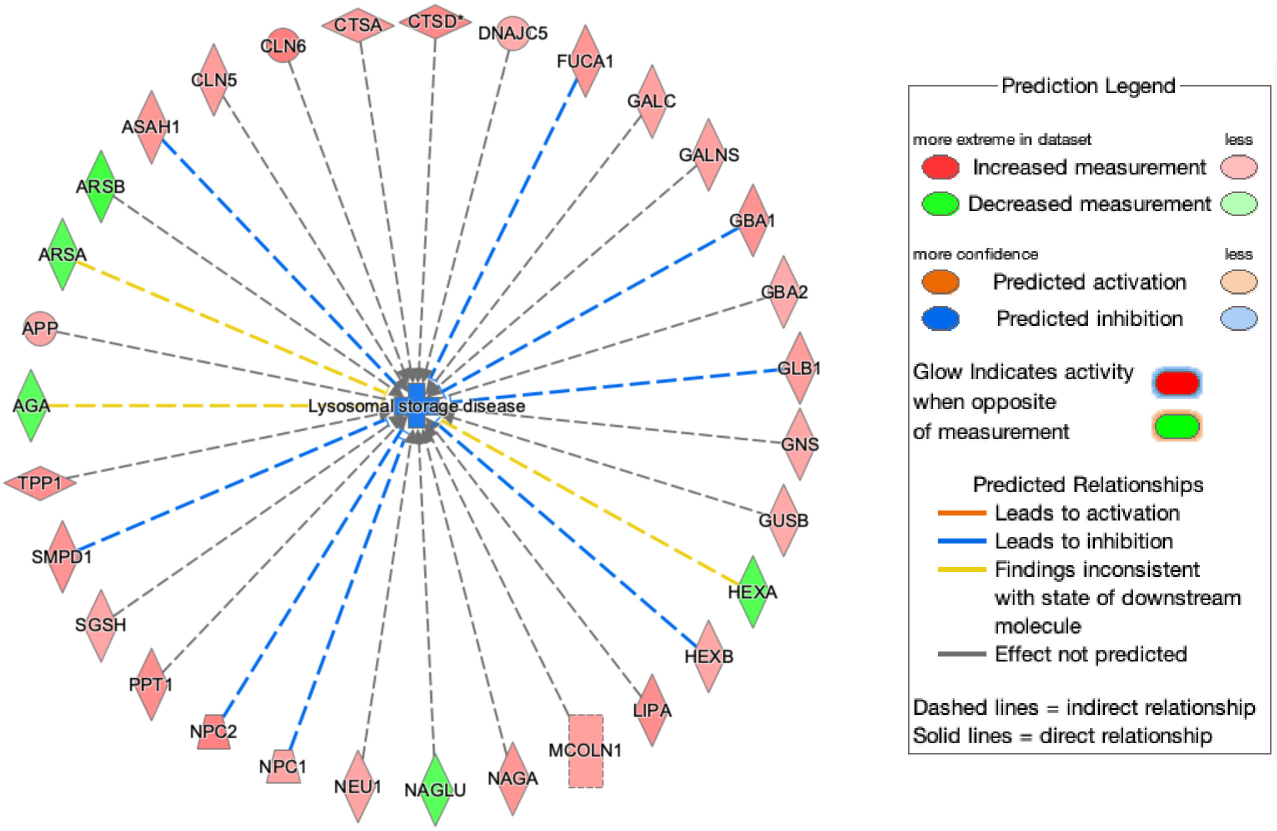

**Supplemental Figure 3**

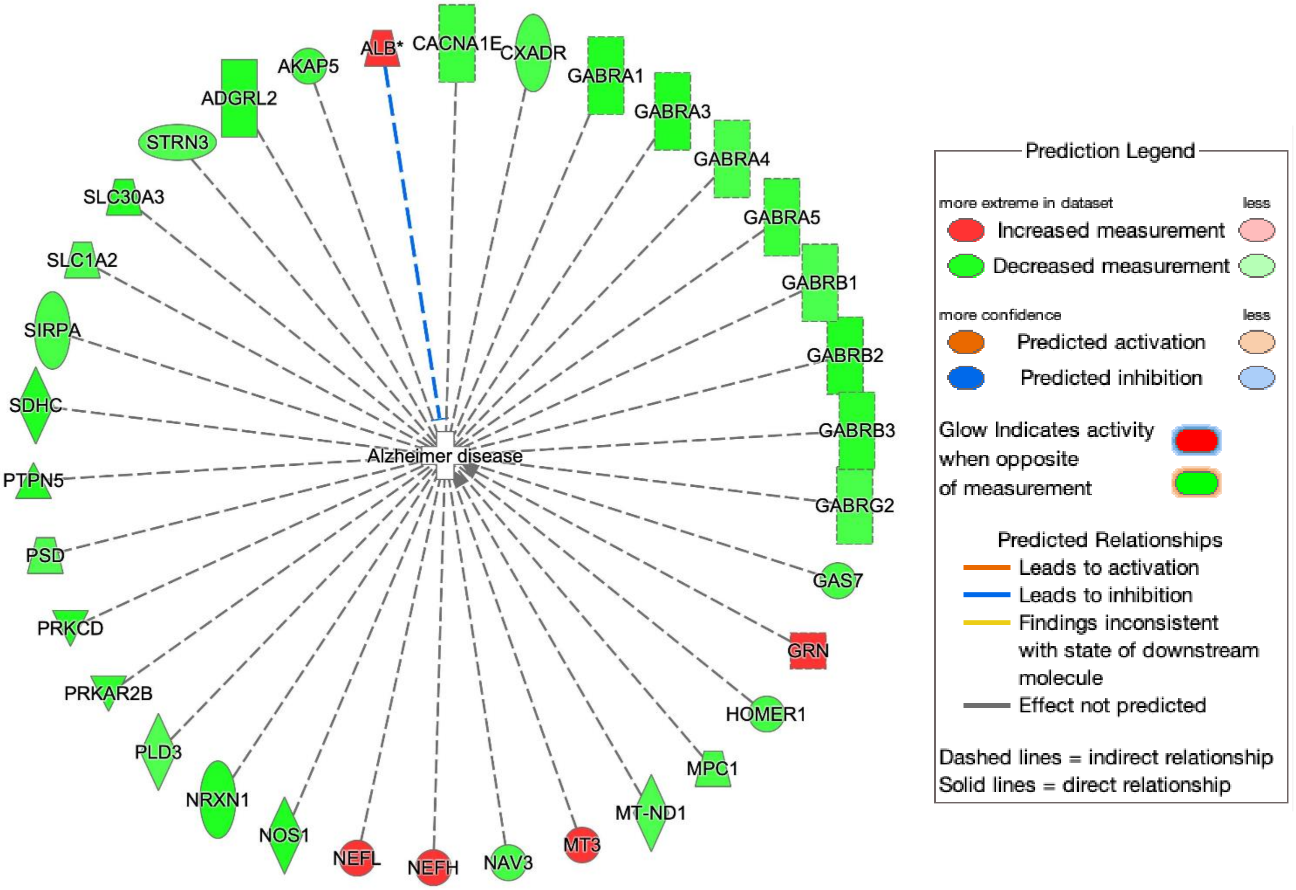

**Supplemental Figure 4**

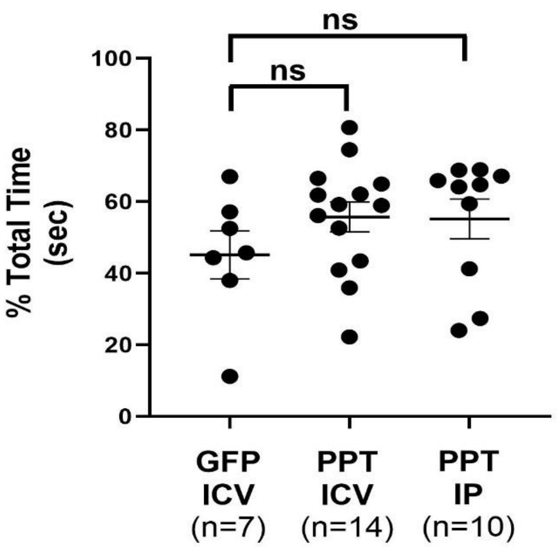

**Supplemental Figure 5**

**A**

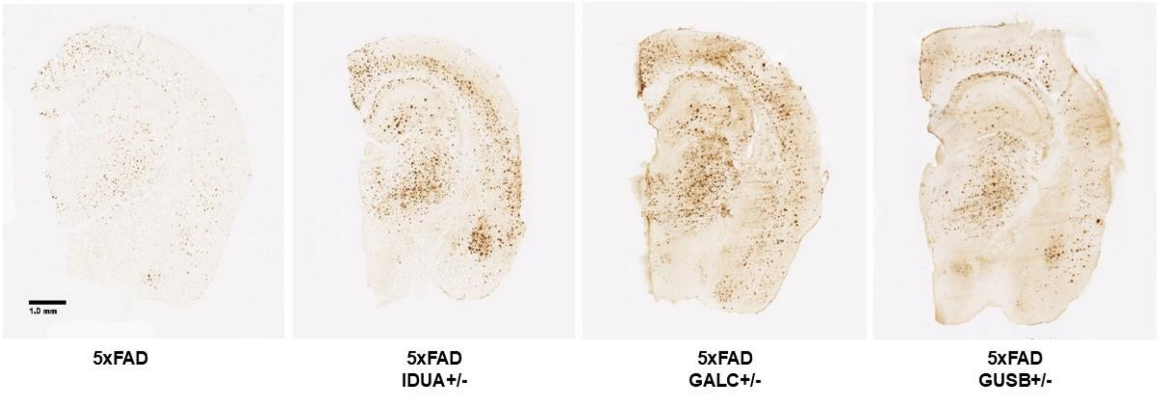


**B**

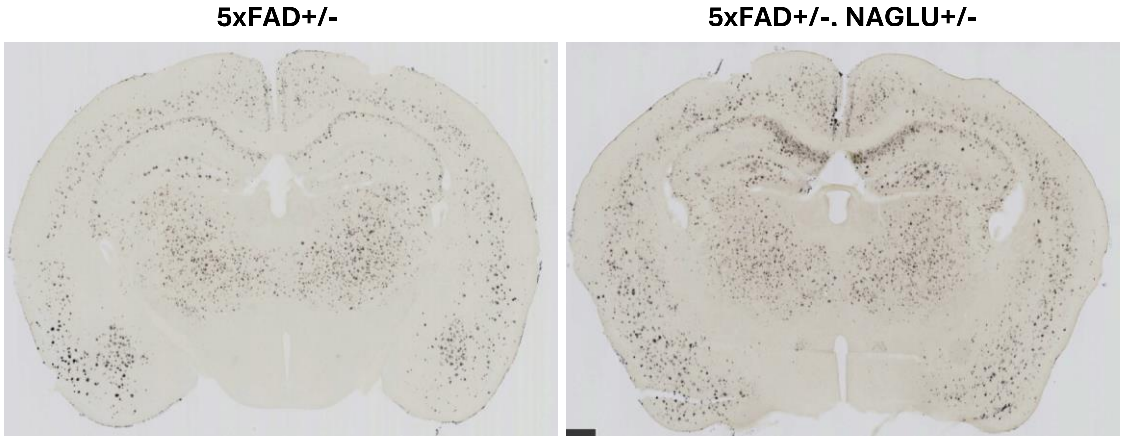


**C**

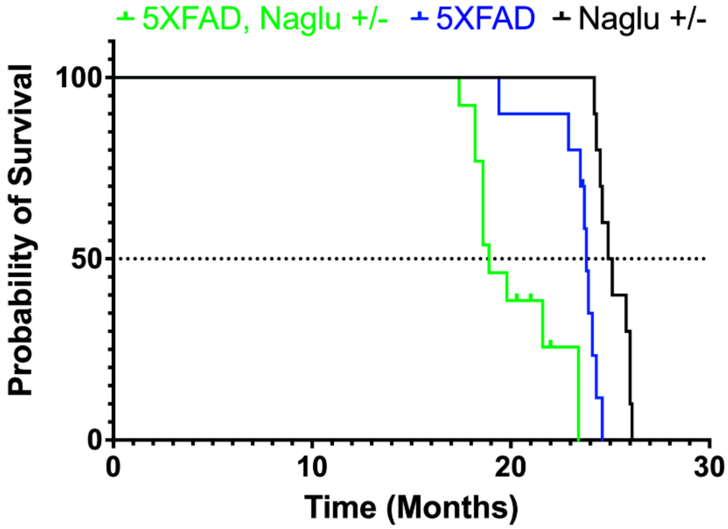
